## Supplementary Information for "FPCountR: Absolute protein quantification using fluorescence measurements"

**Supplementary Note 1. Adaptation of the A280 assay to a plate-based format**

We find that four considerations are required to measure protein concentration by A280 accurately from a Tecan Spark microplate reader. (We assume our findings are translatable to other plate readers.)

**1. Use of UV-clear plastic**. Standard plastic plates used in bacterial growth assays and similar applications are made from polystyrene or polypropylene. Both absorb light under 300 nm very strongly, making them unusable for these assays (1,2). The ideal material for measuring in the UV-range is quartz (3), however plates made of quartz are prohibitively expensive (about £2000 per plate). We find that UV-clear plastic is a good compromise (Supplementary Fig. 4B) and allows for the collection of clean absorbance spectra with clear peaks at 280 nm from the commercially purified protein BSA (Supplementary Fig. 4C).

**2. Path length correction.** As absorbance depends on the path length, and microplate readers necessitate that absorbance readings are carried out from bottom to top, the path length of the sample is not fixed as it is in a cuvette or Nanodrop. Instead, it depends on a number of other factors, such as the volume used, the buffer composition and the temperature. The path length of a microwell may be estimated from its absorbance readings in the infrared range and the k-factor of the same buffer at the same temperature. The k-factor of an aqueous buffer is the observed difference between the absorbance at 975 nm and 900 nm for a path length of 1 cm:

$$k_{buffer|(pl=1)} = A975_{buffer|(pl=1)} - A900_{buffer|(pl=1)} \quad (1)$$

where $k_{buffer|(pl=1)}$ is the k-factor of the buffer, $A975_{buffer|(pl=1)}$ is the absorbance of the buffer at 975 nm, and $A900_{buffer|(pl=1)}$ is the absorbance of the buffer at 900 nm, all taken at a 1 cm pathlength $(pl = 1)$.

Equally,

$$k_{buffer|(pl=w)} = A975_{buffer|(pl=w)} - A900_{buffer|(pl=w)} \quad (2)$$

where all measurements are taken in a microplate well, whose path length in cm, $w$, is unknown. The path length of the liquid in a microplate well is calculated from the ratio of the two k-factors:

$$w = \frac{k_{buffer|(pl=w)}}{k_{buffer|(pl=1)}} \quad (3)$$

It is possible to measure reference k-factors for 1 cm path lengths. However, this requires quartz cuvettes and an appropriate spectrophotometer. More simply, k-factors for many aqueous solutions are available in the literature. We use k-factors from ThermoFisher's reference manual (4), which contains values at 25 $^{o}$C for a range of buffers, as well as how temperature affects them. This data can be viewed in FPCountR with the view_kfactors() function.

In order to apply path length correction to absorbance data, absorbance readings should be taken as absorbance *spectra* from ~200-1000 nm at 1 nm intervals rather than as single readings at 280 nm. This allows the collection of data about both protein concentration (280 nm) and path length (900-975 nm) in the same reading. The function plot_absorbance_spectrum() then uses the raw 975 nm and 900 nm readings to work out the path length of each well, and calculates a path length-corrected value as:

$$A280_{(pl=1)} = \frac{A280_{(pl=w)}}{w} \quad (4)$$

where $A280_{(pl=1)}$ is the sample absorbance at 280 nm at a path length of 1 cm in units of $cm^{-1}$, $A280_{(pl=w)}$ is the sample absorbance at 280 nm at the path length of the well, and $w$ is the path length of the well in cm. Path length correction is carried out for all wavelengths in this way.

Path length-corrected values are then normalised to the mean of the buffer values, to obtain normalised values:

$$normA280_{sample|(pl=1)} = A280_{sample|(pl=1)} - meanA280_{buffer|(pl=1)} \quad (5)$$

where $normA280_{sample|(pl=1)}$ is the normalised absorbance of a sample at a path length of 1 cm, $A280_{sample|(pl=1)}$ is the path length-corrected absorbance of a given sample, and

$meanA280_{buffer|(pl=1)}$ is the mean path length-corrected absorbance value across all the buffer wells. Normalisation is carried out for all wavelengths in this way.

Supplementary Fig. 4D illustrates this process for a dilution series of BSA, and illustrates the fact that large changes in protein concentration can also affect the path length of a solution, presumably through changes in surface tension (Supplementary Fig. 4D, panels i and iii). The absorbance scan processing function plot_absorbance_spectrum() has two other ways of calculating path length (Supplementary Fig. 4D, panel iii). The second uses only the buffer wells for calculation of path length (which can be useful when the FP concentrations are low and the data is noisy) and the third uses the volume only (useful when infrared absorbance readings cannot be taken). The volume method calculates path length using a reference experiment in which a microplate was filled with specified volumes of water (50-300 µl), and the path lengths of each were measured and fitted to a linear model (Supplementary Fig. 4E). A full illustration of the analytical steps of the absorbance scan processing function plot_absorbance_spectrum() in Supplementary Fig. 6 shows how the path length correction and buffer normalisation steps affect an absorbance spectrum.

**3. Removal of common additives**

Initial investigations suggested that the A280 assay overestimated the protein concentrations compared to the microBCA assay. In time, we discovered that two components of the buffer (imidazole and protease inhibitors) both absorb light appreciably at 280 nm (Supplementary Fig. 5), contributing between 0.1-0.4 $cm^{-1}$. This is explained by the ring structures of imidazole (5) and by the fact that commercial protease inhibitors (e.g. Pierce protease inhibitor tablets, EDTA-free, A32955) are a mixture of small molecules and short peptides. Neither component is regularly discussed in the academic or commercial literature as confounders of this assay, therefore a renewed appreciation of this fact might explain why fluorescent proteins, often purified with His tags, might occasionally be thought to possess chromophores that absorb too highly at 280 nm to be quantifiable by their component amino acids. This problem is also unique to the attempt to develop a simple method, as more complex protein purification workflows would inherently include steps following affinity purification that would remove imidazole and protease inhibitors.

**4. Scatter Normalisation**

Once absorbance scans of chemically clean protein samples have been taken and corrected for path length, the concentration of protein can be calculated from the absorbance at 280 nm. Two final analytical factors require consideration. First, as we require data from very low concentrations of protein, beyond the reported limit of sensitivity of this assay in a Nanodrop (100 ng/ul, (6)), the data can be quite noisy. The A280 assay function get_conc_A280() deals with noise by using a loess model fit to the data (Supplementary Fig. 7B) instead of using the raw data points, for identifying the absorbance value at 280 nm. However, knowing the absorbance value at 280 nm (Supplementary Fig. 7D, top panel) is not always sufficient for a precise concentration calculation, as ‘absorbance readings’ are technically readings of the attenuation of light transmittance, not (just) absorption. This is important as attenuation may also be caused by light scatter, which will also decrease the incident light reaching the detector (7,8). The contribution of scatter is removed in a Nanodrop instrument by subtracting the absorbance reading at 340 nm, which is assumed to represent the contribution from scatter at 280 nm (6). The ‘baseline’ normalisation method allows users to remove scatter in this way (Supplementary Fig. 7D, middle panel).

An alternative method for removing scatter uses the mathematical relationship between scatter and wavelength to estimate the scatter at 280 nm more precisely (8). Optical density resulting from Rayleigh scattering is proportional to $1/\lambda^4$ (9). This means that the contribution of scatter is much higher in the UV region than for longer wavelengths, and that at 280 nm it should be twice as high as at 333 nm. The ‘scatter’ normalisation method allows users to remove scatter in this way (Supplementary Fig. 7D, bottom panel). Both normalisation methods allow the user to specify an alternative wavelength – for ‘scatter’, the relationship between the expected scatter at 280 nm and the chosen wavelength is automatically calculated. This can be useful if the FP absorbs around 300-350 nm such as mCherry – recommended settings for mCherry can be found in Supplementary Table 3.

**The ECmax assay**

The procedure for the ECmax assay is very similar to the A280 (Supplementary Fig. 8). Path length normalisation and scatter normalisation are still required, though there is no requirement for UVclear plastic or the removal of imidazole or protease inhibitors. Scatter makes less of a contribution at higher wavelengths, but careful normalisation is still important. Recommended settings for ECmax can be found in Supplementary Table 3.

**Supplementary Note 2. Fluorescence assays are not linear in all buffers**

During the characterisation of calibration protocols experiment illustrated in Fig. 3 and Supplementary Fig. 9, along with protein concentration, fluorescence assays were carried out for each FP to obtain relative fluorescence units to relate to FP molecules in each well. One unexpected observation from this experiment was that fluorescence assays produced textbook linear responses to concentration in buffers containing protease inhibitors, but without these, a steep fluorescence drop was observed, particularly below $10^{12}$ molecules per well (Supplementary Fig. 10A). On first encounter, this result appeared as though it were caused by pipetting error. However, for this experiment, microplates containing dilution series using both buffers were prepared in parallel by the same person, at the same time, using the same equipment, and the result was reproducible across different days and FPs. The lack of linearity could have been due to FP destabilisation in T5N15 buffer, that doesn't contain any stabilising agents. In such case, protease inhibitors could have acted to provide stabilisation in the T5N15+pi buffer. Alternatively, it may have been due to protein adsorption to the plastic at very low concentrations (3), which was apparently avoided by using a buffer containing protease inhibitors. In either case, the observed non-linearity results in an underestimation of the conversion factor in these buffers that scales with gain and resulted in errors of up to 8-fold (Supplementary Fig. 10B), suggesting that the removal of protease inhibitors is inadvisable for an accurate calibration.

**Supplementary Note 3. The use of commercial fluorescent proteins as calibrants**

As described in the main text, we were interested in testing whether our purified FPs gave similar conversion factors as commercially available FPs in order to validate our methods and results, with a focus on GFPs. The result of our efforts to compare commercially available GFPs is provided in Supplementary Table 6.

Unfortunately, we observed several problems with the use of these products as reliable calibrants. First, the majority of the available FPs are either the original “wild-type” *Aequorea victoria* GFP (FPbase: avGFP/1XF1B) – which is almost never used in modern cell biology – avGFP-related but obscure proteins such as alphaGFP (FPbase: B28N7) or Q80R (FPbase 799YV), or commercially-developed proteins such as ‘GFPSpark’ (not on FPbase). Some of these proteins do not have FPbase entries or recorded fluorescence spectra, and others, such as avGFP, have significantly different spectra from the ‘modern’ GFPs like GFPmut3 or sfGFP, absorbing light maximally in the UV range, rather than the blue (488 nm) range. We also found that some FPs were poorly annotated in their datasheets in a way that suggested a lack of quality control (QC; see Supplementary Table 6), and technical support staff occasionally described avGFP as ‘normal’ or ‘standard’ GFP – suggesting an erroneous perception that such GFPs are widely used, or that the identity of the GFP in question is not considered important for most applications. Some suppliers confirmed they do not test their FP batches for fluorescence or carry out QC on their fluorescence activity.

These observations suggested that we would need to validate commercial FPs against our FPs, rather than the other way round. We selected the TurboGFP available from Pierce for testing, and performed calibrations in T5N15pi buffer with the ECmax assay (Supplementary Table 5). Our results confirm that this FP produces similar results to our mGFPmut3 calibrants, with a mean conversion factor of 95.0 % compared to the conversion factor of mGFPmut3. This suggests that users can use commercial FPs to gain approximate conversion factor estimates provided they check for a good match between the properties of their FPs and those available.

Of interest is the fact that in the few weeks since we ordered the Pierce TurboGFP, this product has already been discontinued. For this and the above reasons, we would generally recommend that users make their own calibrants. These are easy to produce, bespoke to the specific FP used in users' own cells and genetic constructs and can be affordably produced in high quantities. Moreover, if users purify FPs and run an SDS-PAGE to ascertain purity, they will have done as much QC as is typically done on commercially purified FPs. As our protocol involves taking an entire absorbance spectrum, this can act as an added QC step to verify FP quality against expected spectra on FPbase. Dilution in protease-inhibitor free buffer and the use of UVclear plates allows users to also check the protein's concentration using two different methods, providing yet another QC step over a commercial protein.

**Supplementary Note 4. Cell count estimates using microsphere calibration**

In the main text, we describe the use of microsphere calibrations to obtain units of molecules per cell (Fig 5). However, we have observed some caveats to the use of microspheres.

We have observed that microsphere calibrations result in errors if they are required to provide conversions *across* OD600 and OD700 data (Supplementary Fig. 13). We suspected that these microspheres may not accurately represent the cellular OD ratio at these two wavelengths. We took absorbance scans of *E. coli* cells and microspheres at different concentrations and analysed the absorbance profiles across the wavelength range 300-800 nm, which confirmed that both particles scatter at shorter wavelengths greater than at longer ones, as expected (10). After normalising to 600 nm wavelength to obtain 'relative OD' values, curves were fitted to each profile. For concentrations represented by approximate OD600/cm values of 0.05-0.5, a non-linear model based on the scattering relationship was fitted:

$$relative\ OD = \frac{k}{(b + \lambda)^4} + a \quad (6)$$

where $\lambda$ is the wavelength in nm, and $a$, $b$ and $k$ are parameters to be fitted. For concentrations represented by OD600/cm values above 1, these were inappropriate and loess models were used instead. The fits are compared in Supplementary Fig. 13A. These show that there are clear differences between the OD700/OD600 ratios for microspheres and cells, particularly for cells in the intermediate concentrations (OD600/cm ≈ 0.5), where for cells the OD700/OD600 ratio is 0.75 and for microspheres it is 0.84, a 12 % difference. (Even larger errors are observed at wavelengths below 600 nm.)

When we compare molecules/cell values obtained from the same experiment, using microsphere calibration values for OD600 vs OD700 measurements (Supplementary Fig. 13B, *left panel*), we find a discrepancy that suggests the OD700 measurements lead to an underestimation of the cell density compared with OD600 measurements, consistent with the observation that the OD700/OD600 ratios are too high. Using a modified OD700 conversion factor that adjusts the OD600 microsphere conversion factor by the average OD700/OD600 ratio obtained from these *E. coli* absorbance scans (0.79), the observed discrepancy disappears (Supplementary Fig. 13B, *right panel*).

For the above reason, we recommend caution using microspheres for calibration of OD for *E. coli* cell counts. As instrument-specific factors effect OD600-specific cell counts, conversion factors for OD600 cannot be precisely predicted, although are likely to approximate empirically obtained values from 1 cm cuvette measurements of ~$1*10^9$ cells/ml for 1 OD600/cm (Supplementary Table 7). Therefore, microsphere calibrations are warranted for these. However, to quantify cells using other wavelengths, an empirical conversion using cellular data should be used, of the form

$$cf_{\lambda=i} = k_{(\lambda=i)} * cf_{\lambda=600} \tag{7}$$

where $cf_{\lambda=i}$ is the conversion factor at wavelength $i$, $k_{(\lambda=i)}$ is the *cellular* OD/OD600 ratio at wavelength $i$ (Supplementary Table 8), and $cf_{\lambda=600}$ is the conversion factor at wavelength 600 nm as derived from a microsphere calibration.

**Supplementary Note 5. FPCount wet lab protocol**

FPCount is a complete protocol for fluorescent protein calibration, consisting of:

1. FP expression and production of cell lysates.
2. FP concentration determination in a microplate reader.
3. FP fluorescence quantification in a microplate reader.

Results can be analysed with the corresponding R package, FPCountR.

This **in-lysate version** of the protocol uses the ECmax protein quantification protocol of **FPs in lysates** and **does not require His-tag purification of the FPs**. Note that it is only suitable for FPs with entries in FPbase (https://www.fpbase.org/).

**CALIBRANT PREPARATION**

**Step 1. Expression**

Set up a 50ml culture for the overnight expression of the calibrant fluorescent protein.

Mix the following in a 200ml flask:

- 50ml culture medium (such as LB Miller)
- 50µl chloramphenicol
- 50µl 20% arabinose (for a final concentration of 0.02%)
- scraping of an expression vector for the calibrant FP in an *E. coli* strain suited to overexpression (e.g. pS381_ara_His-mCherry in *E. coli* BL21 strain)

Incubate the cultures at 30°C overnight with 250 rpm shaking.

**Step 2. Harvesting and Washing**

Harvest the *E. coli* after the overnight culture and wash them to remove media and exchange the buffer to a cell lysis buffer suitable for protein stability. For FPs that express to high levels (e.g. all three FPs from the vectors described in the paper), 20 OD600 units worth of cells contains enough protein for a calibration. For poorly expressing FPs, this can be increased (e.g. to 40 OD600 units of cells), for which users should adjust the relevant quantities below as required.

Prepare the following buffers:

- **Wash buffer = T50N300** (50 mM Tris-Cl pH 7.5, 300 mM NaCl). About 10ml per FP required.
- **Resuspension buffer = T50N300+pi** (T50N300 with protease inhibitors). About 10ml per FP required.
    - Protease inhibitors (EDTA-free, Pierce A32955) are added to the buffer at 1 tablet per 10ml. As these dissolve poorly, it is important to filter sterilise the resultant solutions.
- **Lysis Buffer = T50N300+pi with 100µg/ml lysozyme**. 2ml per FP required (if using 20 OD600 units of cells).
- The following should be available but not pre-prepared into buffers:
    - DNase I (1000U/ml in ddH2O)
    - a stock of $CaCl_2$
    - a stock of $MgCl_2$

Procedure:

- Prechill (for 15min) 1x 50ml falcon tube per FP on ice for cooling the cultures
- Prechill 1x 50ml falcon tubes per FP on ice for sonication (1x falcon tube required per 20 OD600 of cells)
- Prechill a centrifuge with the ability to spin 50ml falcon tubes to 4°C
- Remove culture from incubator. For some FPs it will be clear by eye if expression levels are good.
- Transfer culture to the prechilled falcon tube on ice and cool for 20min. (From here on, cultures and proteins should be kept on ice and spun at 4°C unless otherwise stated.)
- Measure the culture OD600 (use 100µl of culture diluted 1:10 in culture medium) and calculate the OD600 of the original culture.
- Calculate the volume of culture to be aliquoted into the tube for sonication (equivalent of 20 OD600 worth of cells).

Example OD calculation:

| | OD ($cm^{-1}ml^{-1}$) (1:10 dilution) | OD ($cm^{-1}ml^{-1}$) (neat culture) | total OD (OD $ml^{-1}$*50ml) | fraction of culture that is 20 OD | volume (ml) for 20 OD |
|---|---|---|---|---|---|
| mCherry | 0.418 | 4.18 | 209 | 0.096 | 4.78 |

- Add 20 OD to the prechilled tube set aside for aliquoted cultures. (The original cultures can be stored at 4°C for the day or be disposed of.)
- Pellet cells by spinning at ~3,220xg, 10min, 4°C
- Resuspend cells in 5ml Wash Buffer, then add another 5ml Wash buffer

- Pellet cells by spinning at ~3,220xg, 10min, 4°C
- Resuspend in 2ml (for 20 OD) Lysis Buffer (to final concentration of 10 OD/ml).

**Step 3. Lysis.**

Lysis by sonication was found to give the most reliable results. This lysis protocol uses a QSonica Q125 sonicator; settings for other sonicators may vary.

- Stand falcon in small plastic beaker full of ice.
- Lower sonicator tip into the sample.
- Sonicate with settings: amplitude: 50%, pulses: 10s on/10s off, duration: 2min.
- (NB. A 2min duration means 2 minutes of sonication. As this is carried out in pulses of 10s sonication followed by 10s of rest, this takes 4min.)

The lysed solution should go from turbid to clear.

<u>DNase I treatment</u>: DNase treatment is essential if using A280 assay but may not be required for the ECmax assay (this hasn't been tested). Note that DNase I is 31 kDa: similar in size to FPs in a way that may affect yield estimates in an SDS-PAGE, and is sensitive to vortexing.

- To lysates in Lysis Buffer, add:
  - DNase I to 50 U/ml final
  - $CaCl_2$ to 5mM final (13mM ideal for DNase I, <5mM recommended with His resins, (11))

- $MgCl_2$ to 50mM final

- Mix thoroughly
- Incubate for 30min at 4°C

**Step 4. Clarification**

Separate the insoluble fraction from the soluble proteins by centrifugation.

- Transfer lysates from falcons into pre-chilled 1.5ml Eppendorf tubes.
- Spin samples in pre-chilled microfuge at 16,000xg for 30min at 4°C.
- Transfer the supernatant (soluble) fractions to new 1.5ml Eppendorf tubes. Insoluble fractions (1 per FP) may be kept for checking by SDS-PAGE if desired.

**Step 5. Protein concentration (optional)**

In principle, lysates may not always need to be concentrated prior to calibration, but in practice it is recommended to ensure that there is a high enough concentration in the first few dilutions to get accurate protein concentration measurements from the absorbance readings. The total volume of lysate required may in some cases require trial and error, but 1.6ml (equivalent of 16 OD cells) is a good starting point. If 40 OD cells were lysed, up to 4ml lysate can be concentrated at this step. Another good rule of thumb (for green and red FPs at least) is that the FP stock should be concentrated enough to produce visibly colourful solutions.

Using Amicon Ultra 10K columns with 500µl capacity (Merck, UFC5010):

- Add lysate (400µl) to 10K Amicon column (1/n)

- Concentrate by spinning at 14,000xg for 10min at 15$^{o}$C
- Discard flowthrough
- Repeat the above 3 steps as many times as needed. (e.g. n = 4 for 1.6ml lysate)
- Recover resultant sample (~100µl) by eluting into a fresh tube (at 1,000xg for 1min)

**CALIBRATION**

**Step 6. Preparation of FP dilution series**

A calibration plate should be prepared as a serial two-fold dilution series using the same type of 96-well microplate as used for the assays requiring calibration. (If assays are typically conducted using black plates, calibrations should be carried out using the same type of plate.) The requirements for the calibration are 100µl of a concentrated solution of each FP to be calibrated, about 6ml T50N300+pi buffer for each FP, and access to each plate reader to be calibrated. A single dilution series of protein can be used for both the protein assay and the activity assay, and this plate can be stored at 4$^{o}$C if necessary.

**Prepare serial dilution in 1.5ml Eppendorf tubes**

Dilution series are best prepared in 1.5ml Eppendorf tubes as these are easier to handle (and see into) than wells of a deep well plate, which is important for avoiding errors.

A typical dilution series may be prepared as follows.

- Label 11 tubes (e.g. 'mCherry 1' to 'mCherry 11')
- Add 900µl T50N300+pi buffer to tube 1, and 500µl buffer to tubes 2-11.

- Add 100µl concentrated lysate to tube 1 and thoroughly mixed by vortexing and spinning or extensive pipetting.
- 500µl of lysate in tube 1 is removed and mixed (pipetting up and down carefully 8x) into tube 2. This is repeated from tube 2 to tube 3, etc, until tube 11.

**Transfer dilutions into 96-well microplate**

- Arrange plate with each FP occupying a pair of rows (e.g. mCherry in A/B, mGFPmut3 in C/D, etc.). Highest FP concentrations will be placed in the left column (column 1), the lowest near the right (column 11) and buffer in column 12.
- Fill wells in column 12 with 225µl T50N300+pi buffer.
- Transfer 2x 225µl from each FP dilution to the two corresponding wells of the microplate.
- Temporarily cover the plate with a removable lid to prevent it being contaminated (do not seal).

Example plate arrangement (d1 = dilution1 with highest concentration):

| | | 1 | 2 | 3 | 4 | 5 | 6 | 7 | 8 | 9 | 10 | 11 | 12 |
|---|---|---|---|---|---|---|---|---|---|---|---|---|---|
| mCherry | A | d1 | d2 | d3 | d4 | d5 | d6 | d7 | d8 | d9 | d10 | d11 | buffer |
| | B | d1 | d2 | d3 | d4 | d5 | d6 | d7 | d8 | d9 | d10 | d11 | buffer |
| mGFPmut3 | C | d1 | d2 | d3 | d4 | d5 | d6 | d7 | d8 | d9 | d10 | d11 | buffer |
| | D | d1 | d2 | d3 | d4 | d5 | d6 | d7 | d8 | d9 | d10 | d11 | buffer |

**Step 7. The ECmax assay (quantification of FP concentration)**

- Prewarm the microplate reader to the temperature used for standard assays to make sure the calibrations are valid for those temperatures (e.g. 30°C).
- Insert the calibration plate without the lid and equilibrate the calibrants to the plate reader temperature.

- Take an absorbance scan of all filled wells between 200-1000nm. (Wavelengths above 900nm are required for path length correction. Other wavelengths are required for calculation of FP concentration, though UV wavelengths (<300nm) are not always necessary for FPs that absorb at longer wavelengths.)

**Step 8. The fluorescence assay (quantification of FP activity)**

- Remove the plate from the plate reader and seal the plate with a clear plastic seal (e.g. Eppendorf Masterclear real-time PCR film adhesive, 30132947).
- Take fluorescence scans on each plate reader requiring calibration and for each filter set that is used for that FP. Each scan should consist of fluorescence intensity detection at a range of gains (on Tecan Spark gains 40, 50, 60, … 120 are recommended) to allow the relevant scripts to calculate conversion factors for any gain.

**Step 9. Storage of calibrants and calibration plates**

Calibrants should be stored in the fridge, protected from light, where they should be stable for a few weeks after preparation. Calibrant functionality after longer term storage or after freeze-thaw cycles have not yet been tested.

**Step 10. Analysis of calibrations**

The aim of this protocol is to produce conversion factor(s) for any given FP that relate the number of FP molecules to the 'relative' fluorescence units (RFU) observed in a given instrument, with a given filter set, and gain. The previous steps described how to prepare lysates containing FPs to produce calibrants, and how to run the assays that quantify how many FP molecules there are in each well, and the RFU output of each well. This data must now be analysed to obtain the conversion factors required.

The analysis should be carried out using FPCountR, an open-source R package we developed for this purpose. Functions in this package are provided for each analytical step. The parser() functions convert raw plate reader data into tidy data formats. The get_conc_ECmax() function calculates FP concentrations from the absorbance data. The generate_cfs() function obtains conversion factors using concentration and fluorescence data. Finally, process_plate() and calc_percell() functions extract data from microbial growth curves in units of molecules, and molecules per cell.

**Validation steps and controls**

**Expression**

- For some FPs (particularly bright greens and reds), it will be clear by eye if expression levels are high after overnight culture, as the culture will be brightly coloured. If this is not visible in the culture itself, it can become apparent after the first wash step as a brightly coloured cell pellet. For some of these proteins, the absence of colour typically means the expression conditions need optimisation. (Note that some FPs, such as mTagBFP2, will not produce a visible colour.)
- Overnight expression should produce high levels of protein but should not kill the cells: if using a plasmid that results in poor growth (less than 20 OD600 units of cells after overnight culture), use a lower inducer concentration, a lower growth temperature, or some other means to reduce cellular burden.

- Expression should result in adequate to decent yields of soluble protein and minimal aggregated protein. We do not see aggregation with our FPs, although in principle this can be a problem in protein overexpression that would limit FP yields. The presence of aggregates can be checked by SDS-PAGE after the clarification step that separates the soluble and insoluble fraction. The presence of a prominent FP-sized band in the insoluble fraction could indicate protein aggregation. This can often be resolved by using a lower inducer concentration or a lower growth temperature.
- The method has been validated with plasmids that are available on Addgene. These can be used as positive controls if users' own plasmids do not produce enough protein or produce unexpected results.

**Calibrant preparation**

- The lysed solution should go from turbid to clear. If it doesn't, this suggests the lysis was inefficient: repeat the lysis or increase the sonication amplitude or time.
- The clarification step that separates the soluble proteins from the insoluble proteins may be validated by separating the proteins of each fraction (supernatant: soluble proteins; pellet: insoluble proteins) by SDS-PAGE, as shown in Fig. 2C.
- Requirement for concentration: In principle, lysates may not always need to be concentrated prior to calibration, but in practice it is recommended to ensure that there is a high enough concentration in the first few dilutions to get accurate protein concentration measurements from the absorbance readings. The total volume of lysate required may in some cases require trial and error, but 1.6ml (equivalent of 16 OD cells) is a good starting point. If 40 OD cells were lysed, up to 4ml lysate can be concentrated at this step. Another good rule of thumb (for green and red FPs at

least) is that the FP stock should be concentrated enough to produce visibly colourful solutions.

**Protein assay**

- Typical raw data traces from absorbance spectra are provided in Supplementary Fig. 6A. Note the absorbance in the 900-1000nm range used for path length calculation, and the FP-specific absorbance in the intermediate wavelengths. Raw spectra that include wavelengths below 300nm will also contain a characteristic overflow absorbance in the UV range (most plastic microplates absorb in the UV range).
- If the peak is not evident by eye, this can indicate a low starting concentration of protein. After normalisation these usually become apparent, but if they do not, a higher starting concentration of protein is needed.

**Fluorescence assay**

- Visual inspection of raw fluorescence data can usually determine whether the dilutions are reasonably accurate and the replicates reasonably uniform.

**Supplementary Note 6. FPCountR analytical protocol**

Supplementary Note 5 details the protocol for data collection to enable plate reader calibration. We have developed an R package called FPCountR to process this data to (a) determine the instrument/filter set/gain-specific conversion factors for each fluorescent protein, and (b) to combine these conversion factors with *E. coli* time course experiments to determine absolute FP quantities in cells.

All data sets are initially processed by parse() functions that 'tidy' the data into formats easily interpreted by the following functions, with data arranged in columns under meaningful and R-friendly column names and combined with all relevant metadata.

**1. Processing absorbance assay data to retrieve protein concentrations (plot_absorbance_spectrum() and get_conc_ecmax())**

Absorbance assay data (once parsed) is processed by two consecutive functions: plot_absorbance_spectrum() and get_conc_ecmax(). Illustrated examples of the steps in these functions are presented in Supplementary Fig. 6 and 8.

**Normalisation of the absorbance spectra**

Raw absorbance spectra (Supplementary Fig. 6i) for every wavelength (nm) are first corrected for path length = 1cm (Supplementary Fig. 6ii) and secondly normalised to the buffer values by subtraction (Supplementary Fig. 6iii). Both of these steps are described in detail in Supplementary Note 2, point 2.

**Calculation of protein concentrations**

Normalised absorbance values are averaged over replicates, and the resultant mean values of the spectrum are fitted to a Loess model (Supplementary Fig. 8i). In order to convert absorbance values to protein concentration, the FP's molecular weight and extinction coefficient at its excitation maximum (ECmax, $M^{-1}cm^{-1}$) are required.

The molecular weight is obtained using the internal get_mw() function that uses the user-provided protein sequence to calculate its molecular weight in Daltons according to the formula used by Expasy's Compute pi/MW tool (https://web.expasy.org/compute_pi/pi_tool-doc.html):

$$MW = \sum_{i=1}^{n} (aa_i) + w \tag{8}$$

where $MW$ is the molecular weight of a protein in Daltons, $aa_i$ is the average isotopic mass of amino acid $i$ in Daltons, and $w$ is the average isotopic mass of water in Daltons (18.01524 Da). Amino acid masses are obtained from the Expasy database (https://web.expasy.org/findmod/findmod_masses.html#AA). Note that 1 Da is the equivalent of 1 g/mol.

The molar extinction coefficient ($M^{-1}cm^{-1}$) is obtained from FPbase using the internal get_properties() function (Supplementary Fig. 8ii). The molar excitation coefficient obtained from FPbase is converted to a mass excitation coefficient using:

$$mass\ ECmax = \frac{molar\ ECmax}{MW} \tag{9}$$

where $mass\ ECmax$ is the mass extinction coefficient in $(mgml)^{-1}cm^{-1}$, $molar\ ECmax$ is the molar extinction coefficient in $M^{-1}cm^{-1}$ and the $MW$ is the molecular weight in $gmol^{-1}$.

The conversion of absorbance values to concentrations can then be carried out by comparing the absorbance ($cm^{-1}$) value of the Loess fit of each dilution, at the wavelength corresponding to the maximal absorbance wavelength of that FP, as:

$$c_{correction=none} = \frac{Abs_{\lambda=max}}{mass\ ECmax} \tag{10}$$

where $c_{correction=none}$ is the protein concentration in $mgml^{-1}$ (equivalent to the more intuitive unit of $\mu g \mu l^{-1}$) calculated without a correction method (see below), and $Abs_{\lambda=max}$ is the absorbance ($cm^{-1}$) value of the Loess fit at a particular dilution. However, it is usually preferable to correct for light scatter that can contribute to apparent absorbance values before doing this conversion (Supplementary Fig. 8iii; details in Supplementary Note 2, point 4). There are two correction methods. The baseline correction method subtracts the value of the absorbance from another specified wavelength, such as 340 nm:

$$c_{correction=baseline} = \frac{Abs_{\lambda=max} - Abs_{\lambda=340}}{mass\ ECmax} \tag{11}$$

The 'scatter' correction method subtracts the predicted contribution of scatter to light absorbance measurements according to Rayleigh scattering, at a specified wavelength such as 333 nm (details in Supplementary Note 2, point 4). Optical density resulting from Rayleigh scattering is proportional to $1/\lambda^4$ (9), so the expected ratio of light scatter at a the absorbance wavelength versus the normalisation wavelength may be calculated as:

$$scatter\ ratio = \frac{(\lambda_{max})^{-4}}{(\lambda_{norm})^{-4}} \tag{12}$$

where $\lambda_{max}$ is the wavelength of maximal absorbance of the FP and $\lambda_{norm}$ is the normalisation wavelength. For instance, the scatter ratio for mCherry (ECmax at 587 nm) using 333 nm for normalisation would be 0.1. The correction is then calculated as:

$$c_{correction=scatter} = \frac{Abs_{\lambda=max} - scatter\ ratio * Abs_{\lambda=333}}{mass\ ECmax} \quad (13)$$

Finally, a linear model is fitted between dilution and the calculated protein concentration (µgµl$^{-1}$) of each dilution to obtain predicted concentrations in each microplate well (Supplementary Fig. 8iv).

**2. Processing fluorescence assay data to retrieve conversion factors (generate_cfs())**

Fluorescence assay data (once parsed) is fed into generate_cfs(). The generation of conversion factors is carried out by fitting a model that relates protein molecules obtained from get_conc_ecmax() to the relative fluorescence units measured in the fluorescence assay. Our generate_cfs() function is adapted from flopR (12),which uses the method outlined by Beal and colleagues (13,14). Briefly, fluorescence data is normalised by subtracting the average fluorescence of the buffer wells and trimmed of saturated values by removing points that are judged not to be close enough to an accurate two-fold serial dilution (13). Using a specific definition of a saturation threshold of $0.75 * fold\ dilution$ (12) giving a threshold of 1.5 for two-fold dilutions, the function proceeds from low to high concentrations, identifying values in each replicate row that are not at least 1.5-fold higher than the previous value. We modified the flopR function by adding a second saturation check proceeding from high to low concentrations, which serves to clean up the data more effectively. An additional check on low values verifies that these values are distinguishable from the blanks, with the requirement that they be at least 2 blank standard deviations above the mean blank value (14). We also added the normalisation step before the saturation check, unlike in flopR, which allows the preservation of more data points.

Conversion factors were only calculated if at least three data points were left after the above validity checks (14).

Finally, the mean fluorescence value of each dilution is taken, averaging over the replicates, and the conversion factor is determined. For this we use the systematic pipetting error model (equation 7, (14)) to first calculate adjusted protein concentrations due to pipetting error:

$$b_i = p_0 * (1 - \alpha - \beta) * (\alpha + \beta)^{i-1} \tag{14}$$

where $p_0$is the maximum concentration of protein in the highest concentration wells (in molecules per well), $\alpha$ is the dilution factor (defined as $\alpha = \frac{1}{fold\ dilution}$), $\beta$ is the bias due to pipetting error, $i$ is the dilution index and $b_i$ is the expected biased calibrant concentration at the $i$th dilution level. The bias $\beta$ and the conversion factor $cf$ are then simultaneously fit to minimise the sum-squared error over all dilutions (14):

$$\epsilon = \sum_i \left| log\left(\frac{cf * b_i}{nF_i}\right)\right|^2 \tag{15}$$

where $\epsilon$ is the sum squared error of the fit, and $nF_i$ is the normalised fluorescence at the $i$th dilution. (This procedure is repeated for each gain.) The resultant conversion factors are returned by the function to be used in subsequent experimental data processing.

**3. Processing experimental data from *E. coli* timecourse assays to convert arbitrary values to absolute values (process_plate(), calc_fppercell() and calc_fpconc())**

Experimental data from *E. coli* growth and fluorescence assays (once parsed) is processed by the process_plate() function, adapted from flopR with several updates (12).

**Optical density processing:** Optical density (OD) readings from samples are normalised by subtracting the mean of blank wells from the raw OD:

$$OD_{norm} = OD_{raw} - \frac{1}{n}\left(\sum_{i=1}^{n} OD_{blank}\right) \qquad (16)$$

where $OD_{raw}$ is the raw OD, $OD_{blank}$ is the raw OD of blank wells and $n$ is the number of blank wells. These can be further path length-corrected if well volumes are supplied, using

$$OD_{norm|pl=1} = \frac{OD_{norm}}{pl} \qquad (17)$$

where $pl$ is the path length in cm.

**Fluorescence intensity processing:** Fluorescence intensity readings from samples are initially normalised either by subtracting the mean fluorescence of blank wells (af_model = NULL) or by using autofluorescence correction (af_model = “spline”). This uses designated negative controls (samples with cells without fluorescent protein) to generate an OD:fluorescence model that takes into account accumulating autofluorescence often observed at high culture densities (Supplementary method 1 in (12)):

$$flu_{norm=autofluorescence} = flu_{raw} - flu_{neg}(OD_{norm}) \qquad (18)$$

where $flu_{raw}$ is the raw fluorescence of a sample, and $flu_{neg}(OD_{norm})$ is the raw fluorescence of ‘negative control cells’ corresponding to the normalised OD of the sample in question, and $flu_{norm}$ is the normalised fluorescence calculated using autofluorescence correction.

Second, optionally, a correction for cell-based quenching (fluorescence signal attenuation) is applied using the data from Supplementary Fig. 14 which is fitted to the model

$$fc(OD_{norm|pl=1}) = a + \frac{k}{\left(OD_{norm|pl=1} + b\right)^2} \quad (19)$$

where $fc(OD_{norm|pl=1})$ is the fractional fold change in fluorescence given a sample's current normalised OD, $OD_{norm|pl=1}$ is the normalised OD of the cells doing the quenching and $a$, $b$ and $k$ are parameters to be fitted. The correction is applied as:

$$flu_{corrected} = \frac{flu_{norm}}{fc(OD_{norm|pl=1})} \quad (20)$$

where $flu_{corrected}$ is the resultant quenching-corrected fluorescence.

Finally, optionally, OD and fluorescence values are calibrated. Conversion factors ($cf$s) for OD600 or OD700 are generated for each plate reader using the flopR protocol and generate_cfs() function using microsphere calibrations (12). Conversion factors ($cf$s) for FPs are generated from FP calibrants for each plate reader, gain and filter set and using FPCountR's generate_cfs() function as detailed above (equations 14-15). Calibrated values are then calculated as:

$$cells = \frac{OD_{norm}}{cf_{instr|\lambda}} \quad (21)$$

where $cf_{instr|\lambda}$ is the conversion factor (in OD per microsphere particles) of the relevant instrument at the specified wavelength and $cells$ are the cell numbers in particles of equivalent microspheres (PEMS), and

$$FP\ molecules = \frac{flu_{corrected}}{cf_{FP|instr|filter\ set|gain}} \quad (22)$$

where $cf_{FP|instr|filter\ set|gain}$ is the conversion factor (in relative fluorescence units per molecule) of the relevant FP, in the relevant instrument, and at the specified filter set and gain. Finally, $FP\ molecules$ is the protein molecule number in molecules of equivalent FP (MEFP).

Each of the above steps is carried out separately for each time point in a time course assay.

**Obtaining estimates of proteins per cell or intracellular protein concentrations**

Calibrated data may be further processed to protein numbers per cell by dividing the total FP molecules per well by the total cells per well (using the function calc_fppercell()):

$$FP\ molecules\ per\ cell = \frac{FP\ molecules}{cells} \tag{23}$$

Alternately, cellular concentrations can be obtained by dividing FP molecules by the total cellular volume (using the function calc_fpconc()). Estimated cellular volumes may be obtained using:

$$v = OD600_{norm|pl=1} * \frac{g}{10^6} \tag{24}$$

where $OD600_{norm|pl=1}$ is the normalised OD600 (in $cm^{-1}$), $g$ is the OD-specific cellular volume (in $\mu l \cdot OD600^{-1} \cdot cm$), defined in the R function as 3.6 $\mu l \cdot OD600^{-1} \cdot cm$ by default (15), and $v$ is the total cellular volume (in L). (If the OD used to quantify cells in the assay is not OD600, an empirical conversion to OD600 should be performed first (e.g. using data from Supplementary Table 8).) Molar concentrations may then be determined by first calculating the total FP in moles, and secondly by dividing moles of FP by the cellular volume:

$$FP\ moles = \frac{FP\ molecules}{A} \tag{25}$$

where $A$ is Avogrado's number ($6*10^{23}$).

$$FP\ cellular\ concentration = \frac{FP\ moles}{v} \tag{26}$$

where $FP\ cellular\ concentration$ is in Molar units (M).

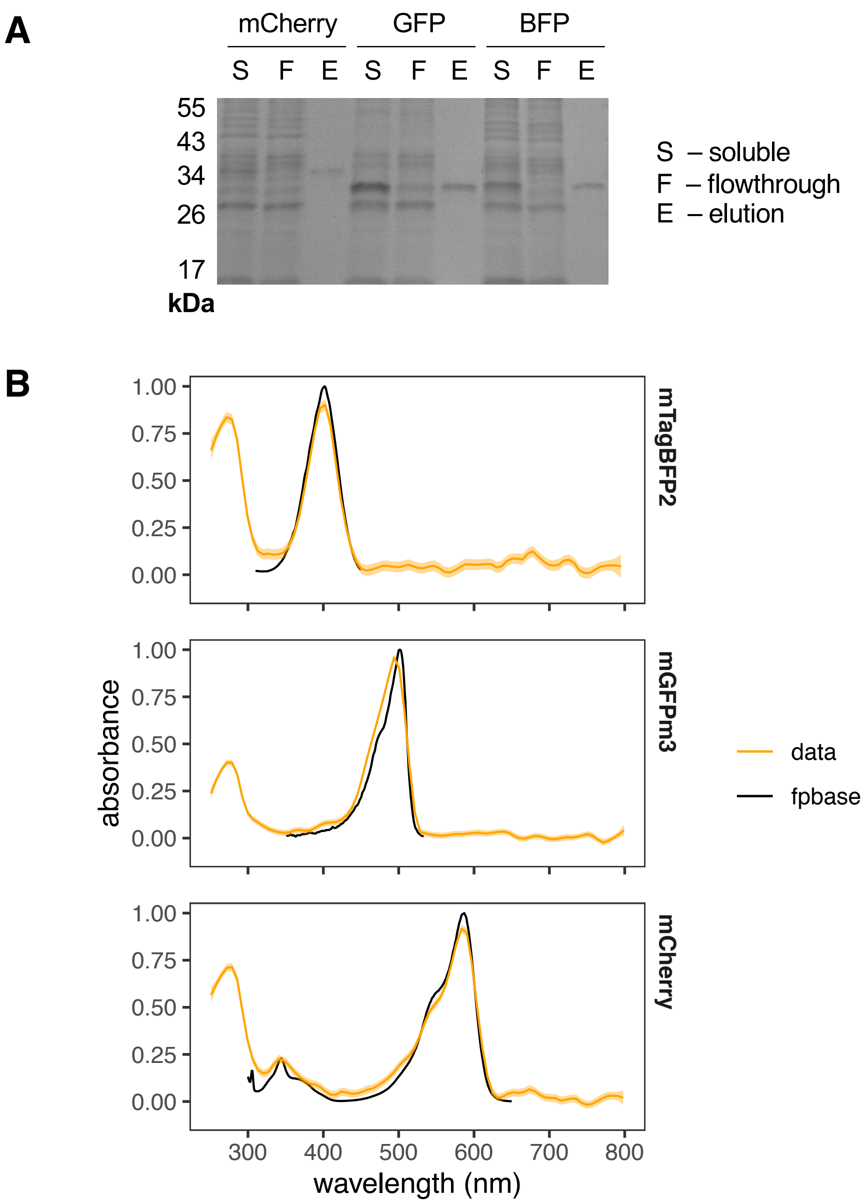


**Supplementary Figure 1. Analysis of purified FPs used in this study.**
**A. SDS-PAGE analysis.** Soluble (S), flowthrough (F) and elution (E) fractions of purifications of mCherry (27.8 kDa), mGFPm3 ('GFP', 28 kDa) and mTagBFP2 ('BFP', 27.8 kDa) were separated on by 12 % SDS-PAGE and stained using Instant Blue. The displayed gel is representative of typical purification results with these FPs. **B. Measured absorbance spectra.** The figure shows obtained absorbance spectra (250-800 nm, normalised such that the highest value = 1) fitted to a loess model with a 95% confidence interval (orange) overlaid with FPbase excitation spectra (black) for each FP. Displayed spectra represent one sample measured in duplicate, that is representative of at least 2 independently purified batches of calibrant.

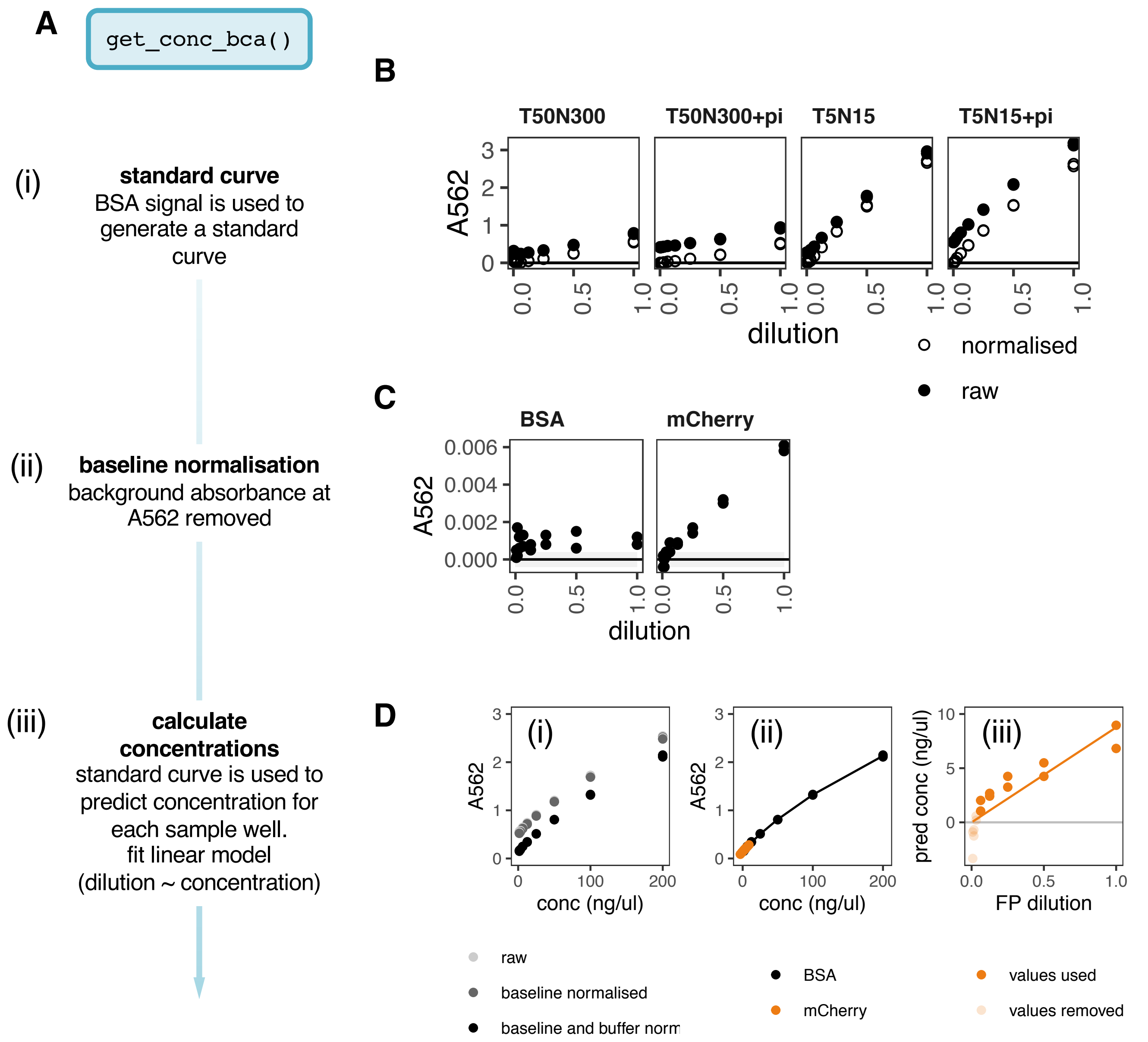


**Supplementary Figure 2. Bicinchoninic (BCA) assay development**
**A. MicroBCA assay protocol.** The original protocol consists of (i) the creation of a (buffer-normalised) standard curve using the provided BSA standard with the assay reagent, and (iii) the comparison of the assay signal of the samples with the standard curve from (i). The amended protocol suitable for FPs, includes an intermediate step (ii) that measures and removes the contribution of the FP to the light absorbance at the assay wavelength (562 nm). **B. Buffer dependence of BSA standard curve.** Serial dilutions of BSA standard were carried out in the indicated Tris buffers (compositions indicated in mM -/+ protease inhibitors, denoted ‘pi’) and measured in duplicate. Plots show the raw (closed circles) and buffer-normalised (open circles) values in duplicate. **C. Baseline absorbance of BSA and mCherry at the assay wavelength, 562 nm.** Baseline absorbance is the inherent absorbance of the protein as opposed to the absorbance due to the BCA reaction. To find the baseline absorbance, it is taken from samples before BCA reagent addition. Plots show the buffer-normalised values of BSA and mCherry absorbance at 562 nm against dilution factor. Grey shading indicates the mean of the buffer ± 2*standard deviations. **D. Example of an analysis with get_conc_bca() using mCherry.** Plots show the raw, baseline-normalised and buffer-normalised BSA standard curve (i), polynomial model fitting to the standard curve and its use to predict FP concentrations (ii), and the predicted concentrations of each FP dilution, with a linear model fit. BSA values are indicated in black, and FP values in orange. Lower transparency points in (iii) indicate values that were removed before fitting the model.

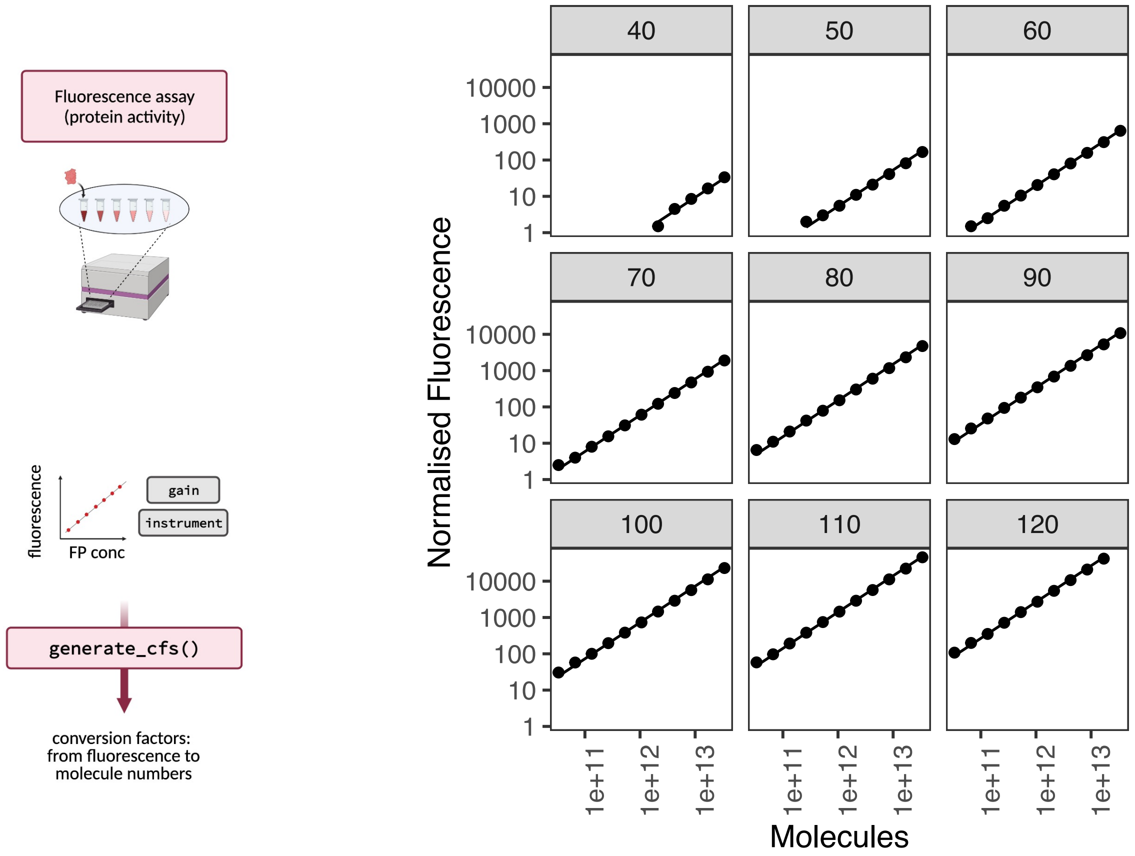


**Supplementary Figure 3. Generation of conversion factors with the generate_cfs() function.**
Visualisation of how the generate_cfs() function works using mCherry. mCherry fluorescence assay values plotted alongside concentration predictions from the microBCA values shown in Supplementary Fig. 2D. The results are separated by instrument gains (40-120).

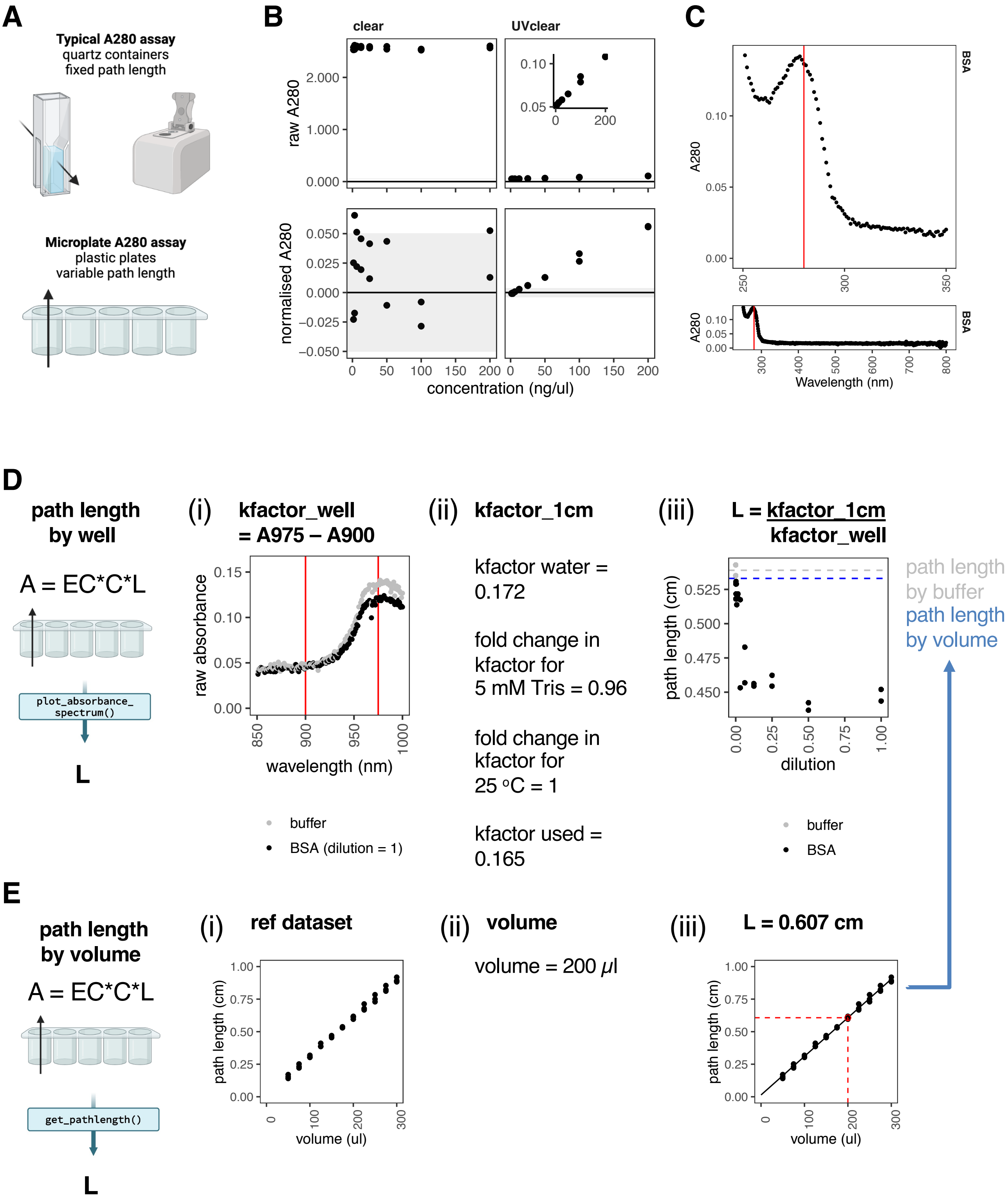
A
Typical A280 assay
quartz containers
fixed path length
Microplate A280 assay
plastic plates
variable path length
B
clear
UVclear
raw A280
normalised A280
concentration (ng/ul)
C
BSA
A280
Wavelength (nm)
D
path length
by well
A = EC*C*L
plot_absorbance_
spectrum()
L
(i) kfactor_well
= A975 – A900
raw absorbance
wavelength (nm)
buffer
BSA (dilution = 1)
(ii) kfactor_1cm
kfactor water =
0.172
fold change in
kfactor for
5 mM Tris = 0.96
fold change in
kfactor for
25 °C = 1
kfactor used =
0.165
(iii) L = kfactor_1cm
kfactor_well
path length (cm)
dilution
buffer
BSA
path length
by buffer
path length
by volume
E
path length
by volume
A = EC*C*L
get_pathlength()
L
(i) ref dataset
path length (cm)
volume (ul)
(ii) volume
volume = 200 μl
(iii) L = 0.607 cm
path length (cm)
volume (ul)

**Supplementary Figure 4. Adaptation of A280 assay to microplates.**

**A. A280 assays** are typically conducted in low-throughput formats such as with cuvettes or in a Nanodrop. To adapt these to microplate formats, challenges such as plate material and path length variation need to be addressed. **B. Use of UVclear plastic allows measurement of A280 values in a plate reader.** A280 values of a BSA dilution series were measured using clear polystyrene (left) and UV-clear plastic (right) microplates. Panels show raw A280 values (*top*) and buffer-normalised A280 values (*bottom*). Inset plot is included to show the relationship between raw A280 values and concentration in UV-clear plates. Grey shading indicates the mean of the blank values ± 2* standard deviations. **C. BSA absorbance spectrum.** An absorbance spectrum of BSA diluted in T50N300 buffer was taken (1 nm resolution, 250-800 nm) and normalised absorbance values are presented for the UV region (*top*) and the entire spectrum (*bottom*). The expected absorbance peak at 280 nm is indicated in red. **D. Illustration of path length estimation and correction.** The function plot_absorbance_spectrum() can estimate the path length on a well by well basis, by (i) the calculation of kfactor_well, the k factor of each well (ii) kfactor_1cm, the kfactor expected if the path length were 1 cm, and (iii) the path length, L. The plot illustrates how this is calculated, using an absorbance spectrum of a serial dilution of BSA in T50N300 buffer. In panel (i) the wavelengths used in the calculation are indicated in red. For clarity, only the highest BSA concentration is shown. Panel (ii) details the calculation of the reference kfactor depending on buffer composition and temperature. Panel (iii) shows the calculated path lengths for the BSA dilution series (data points) vs the path length calculated from the mean of the buffer data (grey line) vs the path length calculated from the known volume only (blue). **E. Path length calculation using volume only.** Data and method for path length estimation using volume. A microplate was filled with a range of volumes of water (n=4), and path lengths for each well were calculated according to the method in D (i). This is the reference dataset used in the package for the calculation of pathlengths using only the known volume (ii). A linear model through the data points is used to calculate the path length for a well with 200 $\mu$l sample - 0.607 cm (iii).

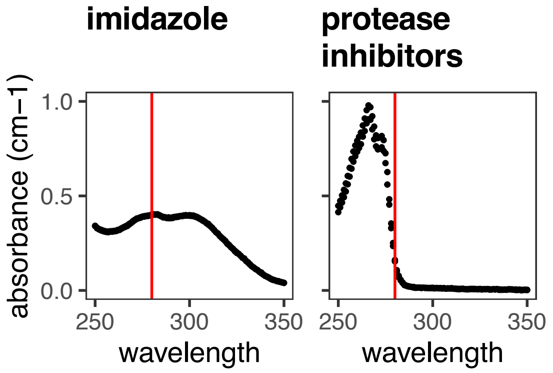


**Supplementary Figure 5. Certain additives are incompatible with A280 measurements.**
Absorbance scans of 150 mM imidazole (*left*) and 1X protease inhibitors (*right*) in T50N300 buffer. The 280 nm wavelength is indicated in red. Points represent duplicate measurements.

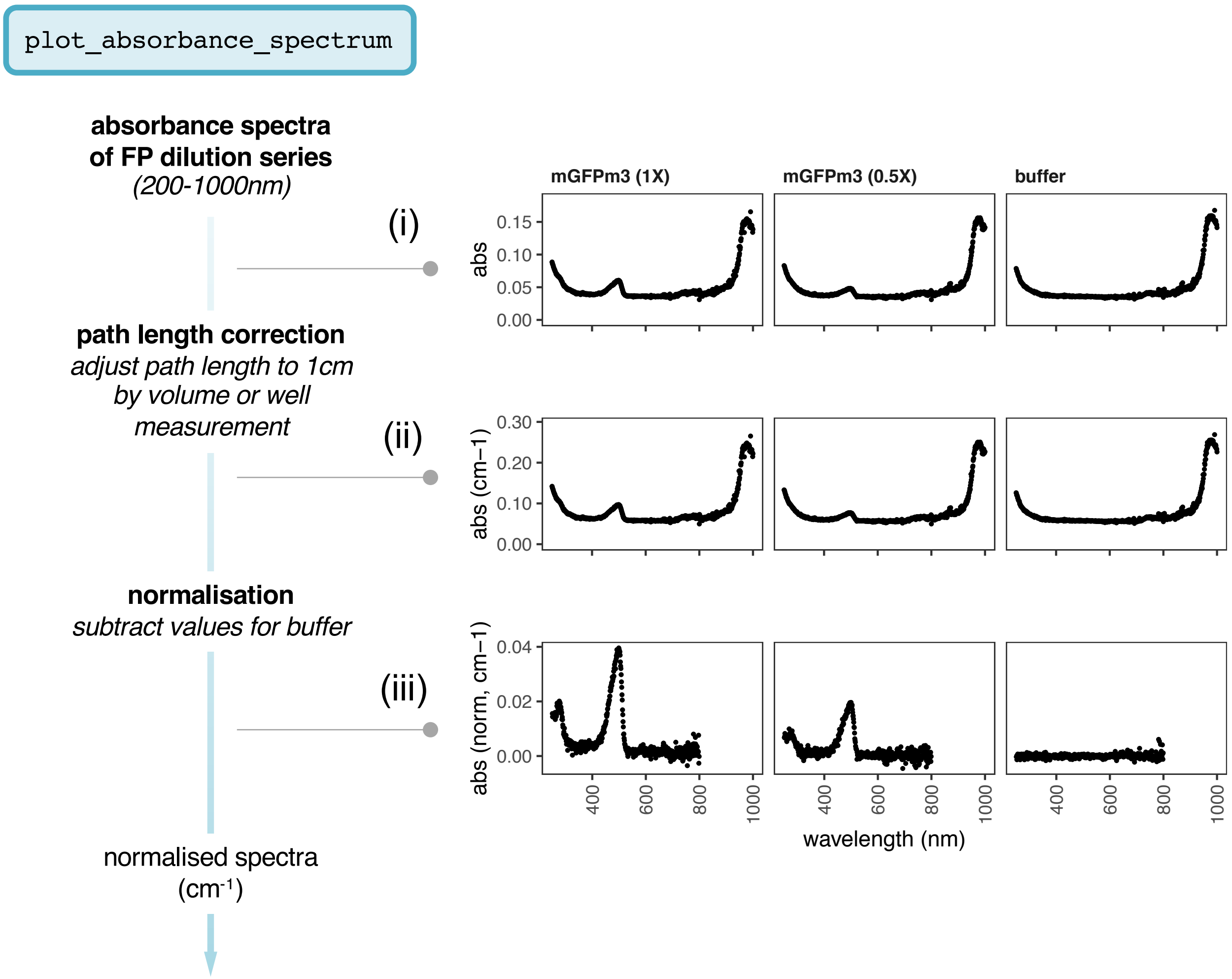


**Supplementary Figure 6. Plotting absorbance spectra with the function plot_absorbance_spectrum().** Illustration of the data processing steps in plot_absorbance_spectrum() on a calibration using mGFPmut3 in T5N15 buffer. Absorbance spectra **(i)** are corrected for path length = 1cm **(ii)** and normalised to the buffer values **(iii)**. For clarity, only two concentrations and the buffer are depicted.

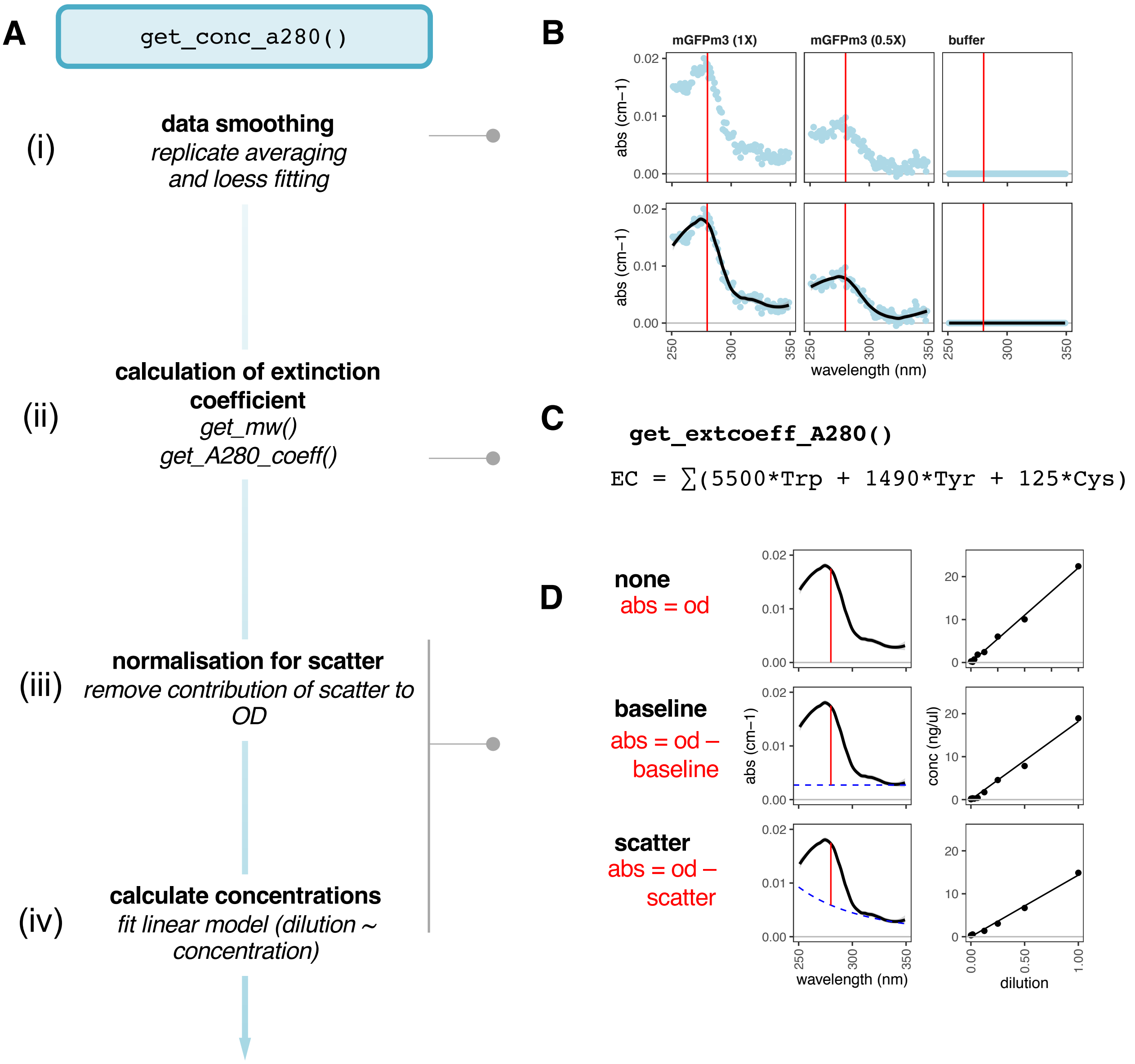


**Supplementary Figure 7. Calculation of protein concentration using the A280 assay.** Illustration of the data processing steps in get_conc_a280() on a calibration using mGFPmut3. **A. Overview.** Normalised values from plot_absorbance_spectrum() are averaged over replicates, and fitted to a loess fit (i). The FP's extinction coefficient at 280 nm (EC280, $M^{-1}cm^{-1}$) is calculated using its protein sequence and molecular weight, using the ProtParam method and the Pace values (ii, see Supplementary Note 1). Measured A280 values are used to calculate the protein concentration at each dilution with or without normalisation (iii). Finally, a linear model is fitted through the predictions for each dilution, and used to determine the protein concentration of the FPs (iv). **B. Loess fitting.** Loess models are fitted to each spectrum from each concentration of FP. Only two FP concentrations and the buffer are shown for clarity. The expected peak at 280 nm is indicated with a red line. **C. Extinction coefficient calculation.** The internal function get_extcoeff_A280() calculates the EC280 of the given FP in $M^{-1}cm^{-1}$ using its protein sequence and the formula shown. This is converted into EC280 in $(mg/ml)^{-1}cm^{-1}$ using the protein's molecular weight. **D. Normalisation for light scatter.** An illustration of how the three scatter normalisation strategies are calculated. No normalisation (*top*) assumes that the optical density at 280 nm is only due to light absorbance by the protein. Baseline normalisation (*middle*) subtracts the OD at 340 nm, and scatter normalisation (*bottom*) subtracts a multiple of the scatter wavelength (333 nm), from the 280nm value to remove light scatter (See Supplementary Note 1 for details.) *Left:* Illustration of normalisation's effect on A280 values used in concentration prediction. Dashed lines represent the contribution of scatter on the optical density predicted by each method that we need to remove. *Right:* Concentration predictions for each method. abs, absorbance, conc, concentration, od, optical density.

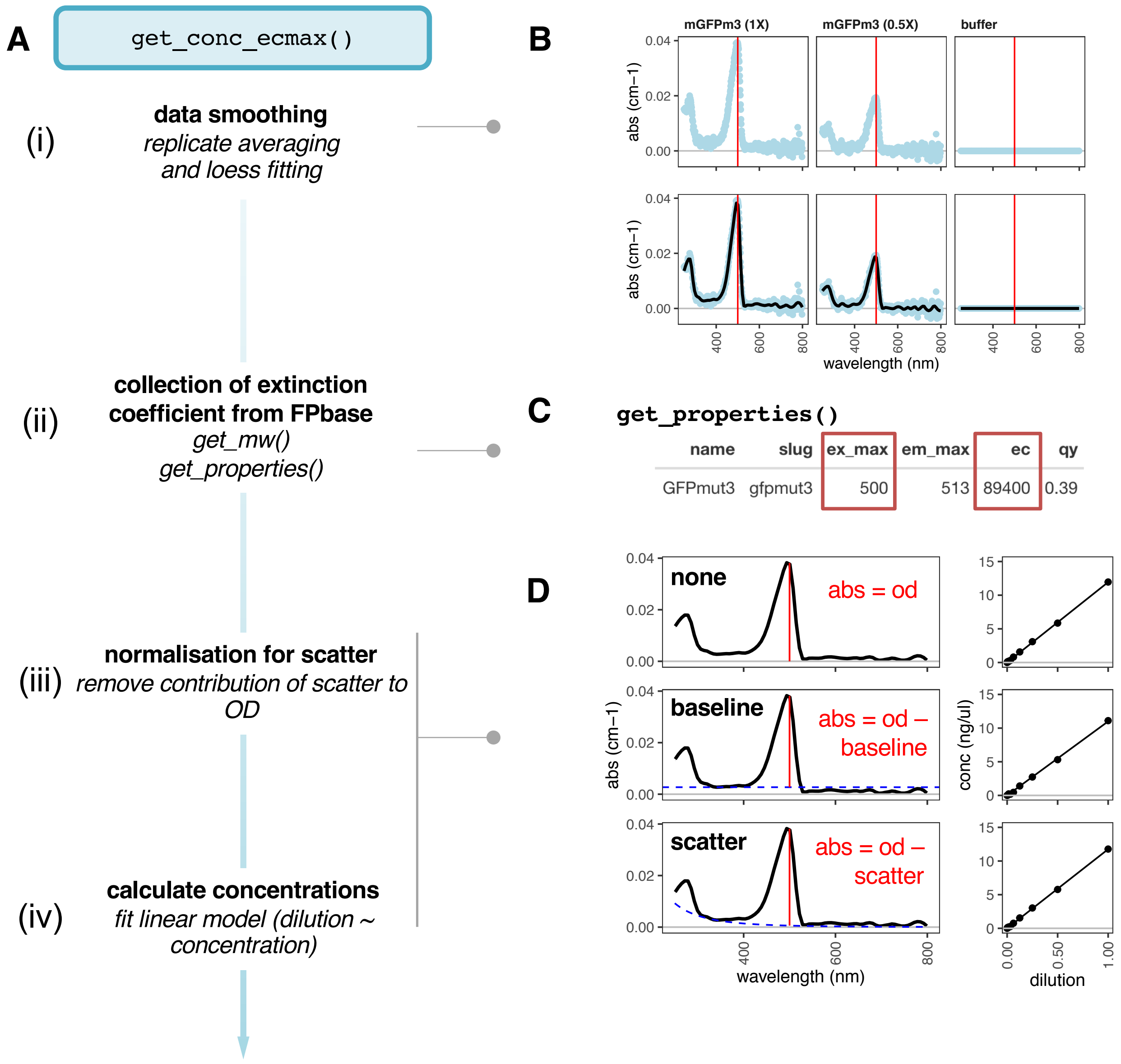


**Supplementary Figure 8. Calculation of protein concentration using the ECmax assay.**
Illustration of the data processing steps in get_conc_ecmax() on a calibration using mGFPmut3.
**A. Overview.** Normalised values from plot_absorbance_spectrum() are averaged over replicates, and fitted to a loess fit (i). The FP's extinction coefficient at its excitation maximum (ECmax, $M^{-1}cm^{-1}$) is fetched from FPbase. Measured ECmax values are used to calculate the protein concentration at each dilution with or without normalisation (iii). Finally, a linear model is fitted through the predictions for each dilution, and used to determine the protein concentration of the FPs (iv). **B. Loess fitting.** Loess models are fitted to each spectrum from each concentration of FP. Only two FP concentrations and the buffer are shown for clarity. The expected peak (500 nm for GFPmut3) is indicated with a red line. **C. Extinction coefficient retrieval.** The internal function get_properties() retrieves key properties of the given FP such as its excitation maximum (here, 500 nm) and its ECmax (here, 89,400) in $M^{-1}cm^{-1}$. This is converted into ECmax in $(mg/ml)^{-1}cm^{-1}$ using the protein's molecular weight. **D. Normalisation for light scatter.** An illustration of how the three scatter normalisation strategies are calculated. No normalisation (*top*) assumes that the optical density at the ECmax wavelength is only due to light absorbance by the protein. Baseline normalisation (*middle*) subtracts the OD at 340 nm, and scatter normalisation (*bottom*) subtracts a multiple of the scatter wavelength (333 nm), from the ECmax wavelength value to remove light scatter (See Supplementary Note 1 for details.) *Left:* Illustration of normalisation's effect on ECmax values used in concentration prediction. Dashed lines represent the contribution of scatter on the optical density predicted by each method that we need to remove. *Right:* Concentration predictions for each method. abs, absorbance, conc, concentration, od, optical density.

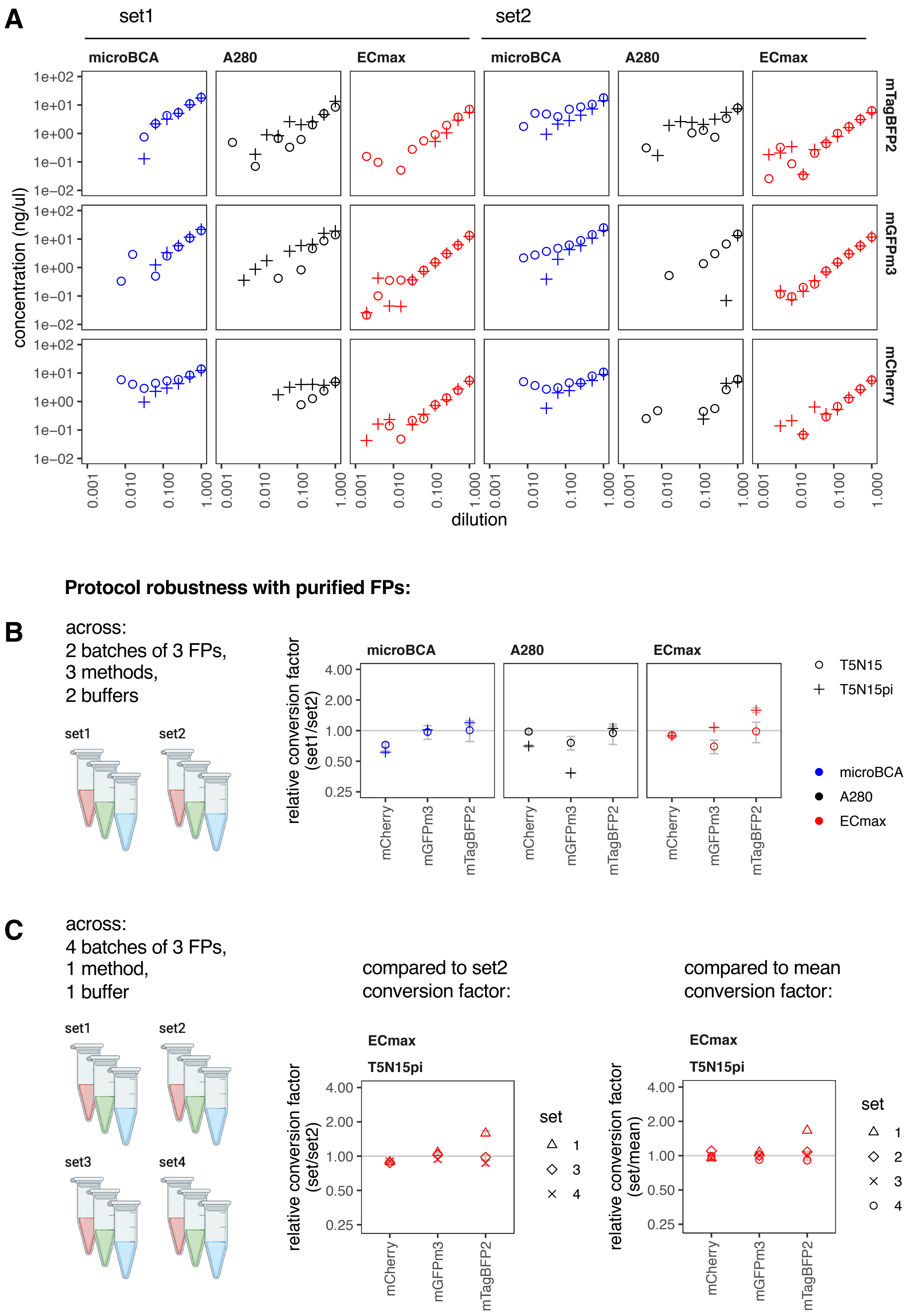

A
set1
set2
microBCA
A280
ECmax
mTagBFP2
mGFPm3
mCherry
concentration (ng/ul)
dilution
Protocol robustness with purified FPs:
B
across:
2 batches of 3 FPs,
3 methods,
2 buffers
set1
set2
relative conversion factor (set1/set2)
T5N15
T5N15pi
microBCA
A280
ECmax
C
across:
4 batches of 3 FPs,
1 method,
1 buffer
set1
set2
set3
set4
compared to set2 conversion factor:
compared to mean conversion factor:
ECmax
T5N15pi
relative conversion factor (set/set2)
relative conversion factor (set/mean)
set
mCherry
mGFPm3
mTagBFP2

**Supplementary Figure 9. Effect of protein assay and buffer on concentration and conversion factor estimation of three FPs.**

**A. Full data from the systematic comparison of three FPs across two buffers, three assays and two purifications.** Plots show the measured concentrations of each FP dilution (FPs in rows) across two purifications (set1 and set2) with the three assays (microBCA in blue, A280 in black, ECmax in red), across two buffers (open circles, T5N15; crosses, T5N15 with protease inhibitors). Each serial dilution was a 2-fold dilution, where 11 dilutions were prepared and measured in duplicate. Points represent the mean of the duplicate values. All 11 dilutions were measured with the A280 and ECmax assays but only the top 8 dilutions were tested with the microBCA assay. The values for the A280 and ECmax assays were normalised for scatter as indicated in Supplementary Table 2. Any missing data points had concentrations recorded as being below 0.01 ng/$\mu$l. **B. Comparison of conversion factor prediction across batches.** Conversion factors (cf) across batches (set1 vs set2) were compared by calculating the fold difference at each gain. Plots display mean fold differences across the gains and error bars represent the standard deviations. Colours and shapes are identical to (A). **C. Comparison of conversion factor prediction across further independent repeats using ECmax assay and T5N15pi buffer.** Showing conversion factors across the two existing sets of purifications in A-B (sets1-2), plus two further sets (sets 3-4) compared to conversion factor of set 2 (*left*) or the average conversion factor (*right*). Note that the set 1 value for mTagBFP2 was excluded from the calculation of the average cf (*right*).

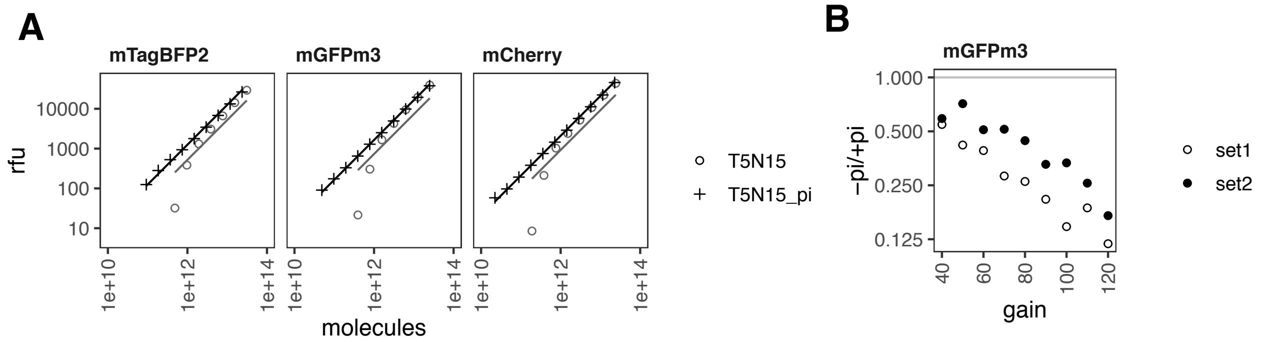


**Supplementary Figure 10. Effect of buffer on fluorescence assays and conversion factor estimations.** Fluorescence assay data from the comparative experiment presented in Figure 3 and Supplementary Figure 9. **A. Effect of buffer on fluorescence assays.** Combining the protein concentration predictions from the ECmax assay (x axis) with relative fluorescence units of the instrument to be calibrated from the fluorescence assay, conversion factors may be calculated by fitting a linear model of relative fluorescence units vs. molecules of protein. Points represent the mean of duplicate values at each concentration, and the line represents the fit found by the generate_cfs function. Patterns of linearity evident in these plots are representative of all fluorescence assays carried out in these two buffers. **B. Effect of buffer on conversion factor calculation.** Conversion factors were calculated for each protein, set and buffer using the modelled fits between relative fluorescence units and protein concentration from the ECmax assay, as in A. The fold difference in conversion factor prediction in buffers without (-) and with (+) protease inhibitors (pi) are displayed for mGFPm3, across both batches (set1 and set2) of protein. Each point therefore represents one value for each FP batch.

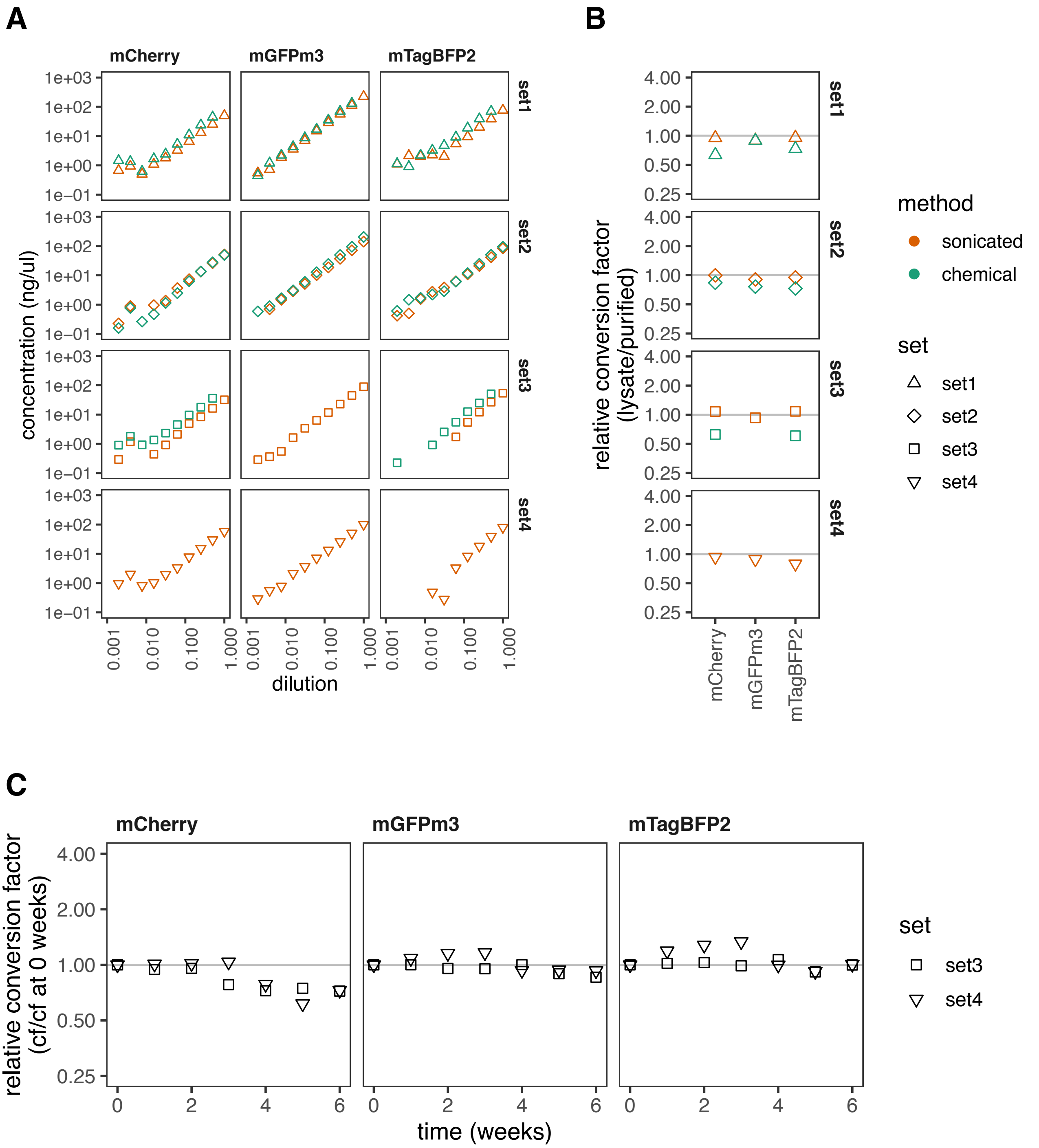


**Supplementary Figure 11. Performance of FPs in lysate as calibrants. A-B. Full data from independent repeats of the lysate protocols shown in Fig. 3E-F.** Plots show the measured concentrations (A) and relative conversion factors compared to purified FPs (B), across up to 4 sets of experiments using lysis by sonication (orange) or chemical lysis (green) using the ECmax assay and T50N300+pi buffer. **C. Stability of two sets of sonicated lysates over a period of storage at 4ºC.** Two of the batches of calibrants prepared for A-B were stored at 4ºC and remeasured over a period of six weeks. Plots show their measured conversion factors relative to the conversion factors measured at 0 weeks.

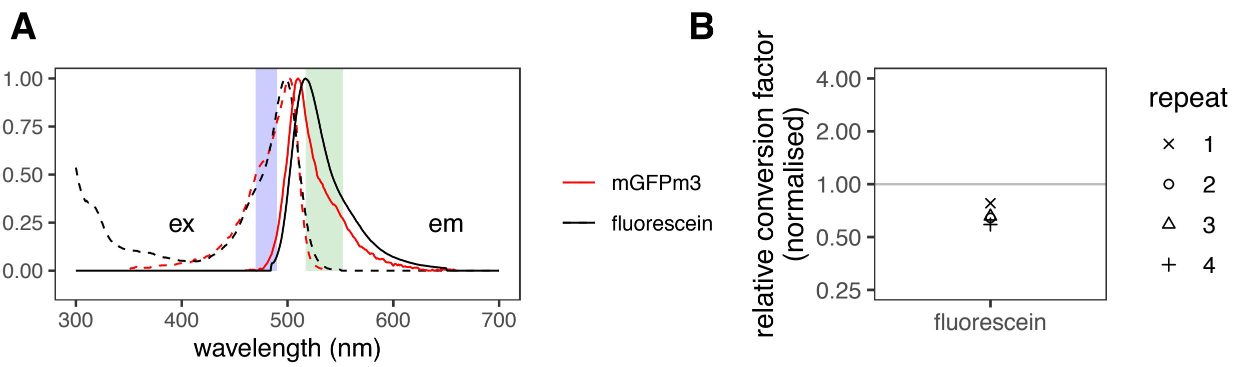


**Supplementary Figure 12. Comparison of the conversion factors of mGFPm3 and fluorescein. A. Fluorescence spectra comparison.** Excitation (ex, dashed lines) and emission (em, solid lines) spectra of mGFPm3 (red) and fluorescein (black). The excitation (blue-) and emission (green-) shaded areas represent the bandpass filter wavelengths of our GFP-specific filter sets (respectively, 480/20 and 535/25 nm). **B. Relative conversion factors.** A serial dilution of known concentrations of fluorescein was prepared and fluorescence intensity values were taken with the same instrument, filter set and gain as the mGFPm3 calibrations done previously. Calculated conversion factors were adjusted for the expected difference in brightness and excitation/emission in the 'GFP' filters. Each point represents one independent experiment taken from a serial dilution prepared in duplicate.

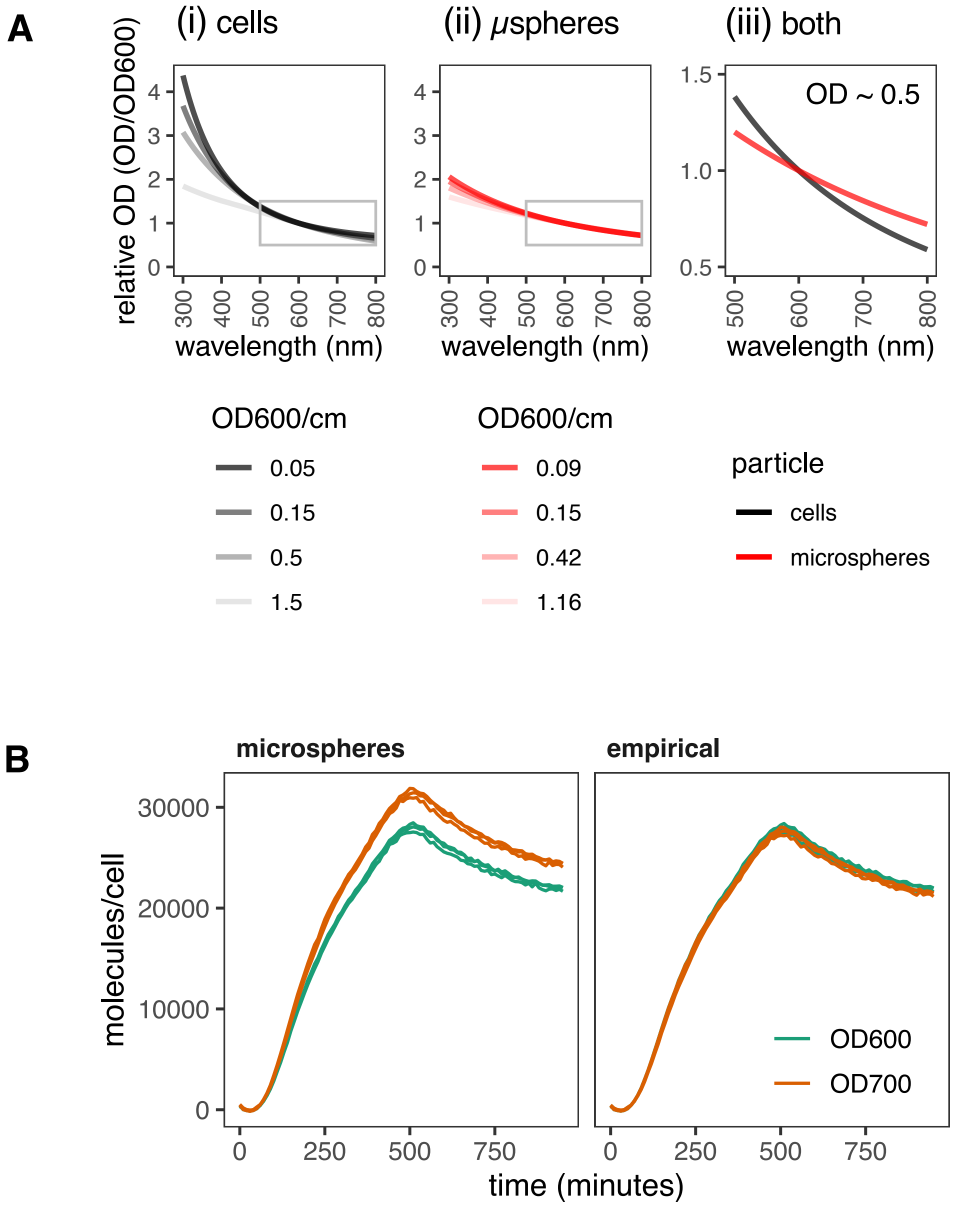


**Supplementary Figure 13. Validity of microspheres as particle number calibrants across wavelengths. A. Absorbance spectra comparisons.** Absorbance scans of *E. coli* cells (i) and microspheres (ii) were collected at a range of concentrations, and plotted as normalised to OD600 to allow comparison between different concentrations. Panels show model fits only for clarity. The grey box illustrates the area used for panel (iii) zoom. Panel (iii) shows a direct comparison between cells and microspheres at an approximate OD of 0.5 (cells: 0.5, microspheres: 0.42). **B. Performance of two methods of cell number calibration.** DH10B cells transformed with pS361_ara_mTagBFP2 plasmid were induced with 0.3% arabinose and followed with OD600, OD700 and fluorescence measurements. Proteins per cell were calculated using calibrated OD/particle values from either microsphere calibration for both OD600 and OD700 channels (*left panel*), or only for the OD600 channel *(right)*. In the latter case, the OD700 conversion factor was calculated as OD600 conversion factor * 0.79 using the data in (A) and Supplementary Table 5. Data was collected and is displayed in triplicate.

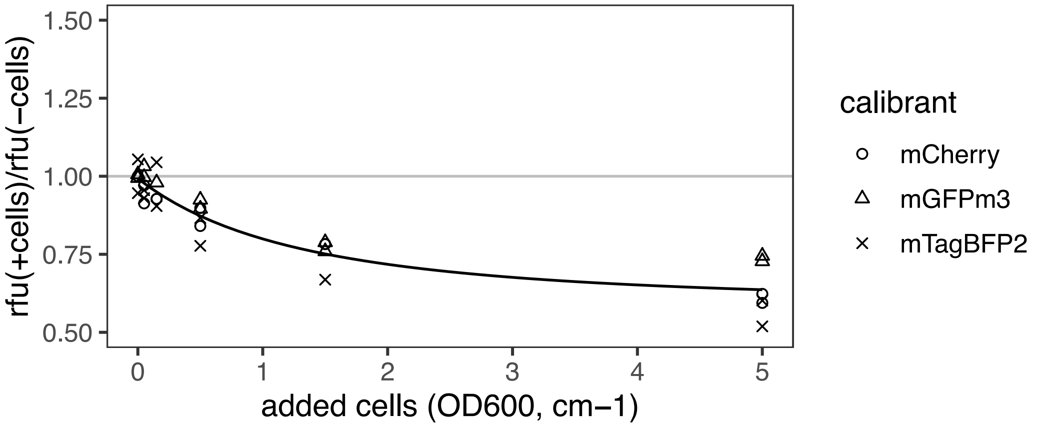


**Supplementary Figure 14. Quenching of fluorescence by *E. coli* cells.**
The effect of *E. coli* DH10B cells on the measured fluorescence of three purified FPs was quantified. To an equal amount of each FP in T5N15 buffer containing protease inhibitors, increasing concentrations of cells were added and mixed. OD and fluorescence was recorded. Raw fluorescence was corrected for cellular autofluorescence, before cellular quenching was calculated as the relative fluorescence units (rfu) of samples containing FP with cells, compared with samples containing only FP. Data was collected in duplicates and a model of form $y = k / (x+b)^2 + a$ was fitted through all data points, as shown.

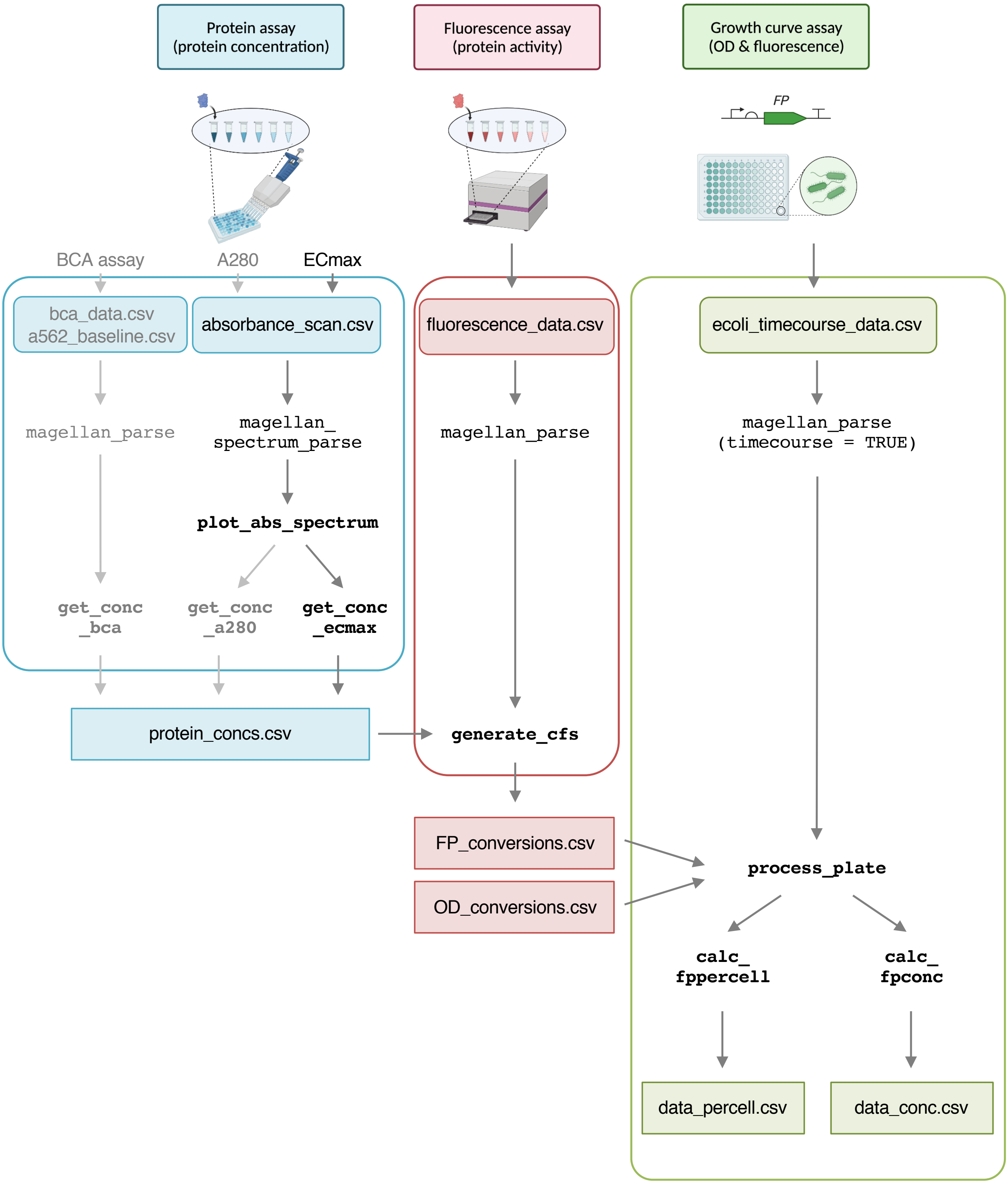
Protein assay
(protein concentration)
Fluorescence assay
(protein activity)
Growth curve assay
(OD & fluorescence)
FP
BCA assay
A280
ECmax
bca_data.csv
a562_baseline.csv
absorbance_scan.csv
fluorescence_data.csv
ecoli_timecourse_data.csv
magellan_parse
magellan_
spectrum_parse
magellan_parse
magellan_parse
(timecourse = TRUE)
plot_abs_spectrum
get_conc
_bca
get_conc
_a280
get_conc
_ecmax
protein_concs.csv
generate_cfs
FP_conversions.csv
OD_conversions.csv
process_plate
calc_
fppercell
calc_
fpconc
data_percell.csv
data_conc.csv

**Supplementary Figure 15. A full version of Figure 1's overview of the FPCountR package's functions.**
Raw data files exported by a microplate reader are represented in curved boxes, R functions are represented in `courier font`, and data files produced by the analysis are presented in rectangular boxes. Protein assay data for the BCA assay is processed by the **get_conc_bca** function (Supplementary Fig. 2), while A280 and ECmax data from absorbance scans are processed by sequentially by the **plot_absorbance_spectrum** function (Supplementary Fig. 6) and the **get_conc_a280** function (Supplementary Fig. 7) or **get_conc_ecmax** function (Supplementary Fig. 8), respectively. These create a file containing protein concentrations. Together with the fluorescence data, the protein concentration file is used to find conversion factors using **generate_cfs** (Supplementary Fig. 3). Files containing fluorescent protein and OD conversion factors are then fed into each analysis of experimental data, first using **process_plate** that normalises and calibrates the data, as well as removing cellular autofluorescence and cellular fluorescence quenching (Fig. 4-5), then either by **calc_fppercell** that provides 'molecules per cell' values, or by **calc_fpconc** that provides molar concentrations (Fig. 5). (Each set of raw data also requires a parser function that formats the raw data as it is exported from the plate reader.)

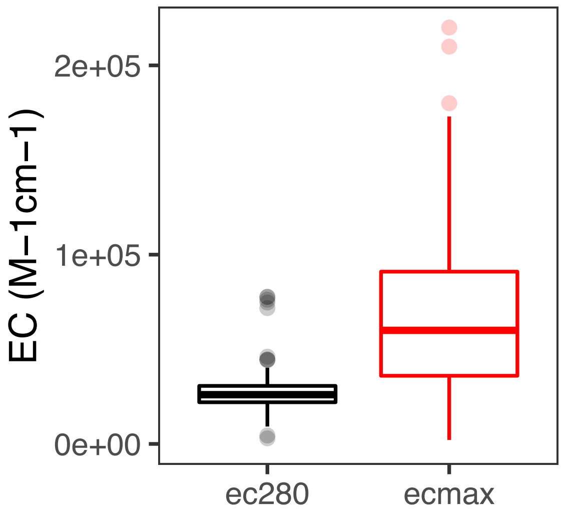


**Supplementary Figure 16. Distribution of extinction coefficients for all FPs on FPbase.**
Box plot shows the distribution of all fluorescent proteins' extinction coefficients at 280 nm (EC280) and at their maximal excitation wavelength (ECmax) in $M^{-1}cm^{-1}$. The EC280 values were calculated from the protein's primary sequence and ECmax was taken from its list of properties. Plots are displayed to indicate the median and quartiles, outliers (more than 1.5*the interquartile range above the third, or below the first, quartile) are shown as individual points.

**Supplementary Tables**

**Supplementary Table 1. Predicted concentrations of all FPs in protein assay comparison experiment from Figure 3/Supplementary Fig. 9.**
Columns1-6: Concentration calculations for each FP set, by each method, over two buffers. Each value represents the concentration prediction for the highest concentration in each serial dilution (where dilution = 1), where each serial dilution was measured in duplicate (see Supplementary Fig. 2, 7 and 8 for details). Columns 7-9: Prediction error compared to the A280 reference (the A280 value of the T5N15 buffer for that sample). Errors were calculated as the assay value divided by the reference value.

**Supplementary Table 2. Root mean square error statistics for protein quantification assay data.** [1]Mean RMSE across all proteins and data sets in Supplementary Figure 9A. [2]The 'pi' notation indicates the inclusion of protease inhibitors.

**Supplementary Table 3. Recommended scatter normalisation settings for each FP.**
Both A280 and ECmax assays require scatter normalisation (correction for the contribution of scatter to their absorbance) by 'baseline' or 'scatter' methods (see Supplementary Fig. 7-8). Such normalisation uses the absorbance at a value other than the wavelength used for concentration prediction (not 280 nm or not the FP's ECmax wavelength) in order to estimate the contribution of scatter to the absorbance at the wavelength used for concentration prediction. This table details the wavelengths recommended for the scatter normalisation of mTagBFP2, mGFPmut3 and mCherry. The units for each column is in nm. The default settings are 340 nm for baseline normalisation and 333 nm for scatter normalisation. Where settings are different from the defaults, they are shown in bold. The columns "correction preferred" indicate which of the correction methods is ultimately recommended. These were also the settings used for the data in Supplementary Fig. 9.

**Supplementary Table 4. Extinction coefficients of FPs at 280 and maximal excitation.**
Extinction coefficients at 280 nm and the excitation maximal wavelength (in $M^{-1}cm^{-1}$) of the three FPs used in this paper and the average of all FPs according to FPbase. The final column shows the fold difference between the ECmax and the EC280 for each row (rounded to 2 decimal places).

**Supplementary Table 5. Relative conversion factors.** [1]Conversion factors (relative fluorescence units/molecule number) from purified, sonicated lysate and chemical lysate calibrants are all shown as relative to the mean of the purified equivalent fluorescent proteins (purified mCherry values were compared to the mean of all <u>purified</u> mCherry values, and sonicated mCherry lysate values were also compared to the mean of <u>purified</u> mCherry values, etc). [2]One anomalous value (mTagBFP2 purification set1) was removed from the calculation of the mean as described in the text. [3]Standard deviation. [4]Coefficient of variation. [4]Sets here refer to independent replicates. [5]TurboGFP and fluorescein are shown as compared to the purified mGFPmut3 samples.

**Supplementary Table 6. Comparison of commercially available GFPs.** The properties of purified GFPs available as of May 2022 listed in order of suitability as calibrants for calibration of the mGFPmut3 protein used in this work. The left of the table compares the

GFPs in terms of their identity with FPbase-recorded protein sequences and fluorescence properties, and the right in terms of the purification and QC procedures of the supplied proteins. [1]Slug – FPbase slug; [2]ID – FPbase ID; [3]EC – extinction coefficient according to FPbase; [4]QY – quantum yield according to FPbase; [5]Brightness – brightness according to FPbase; [6]Ex max – maximum emission wavelength; [7]Em max – maximum excitation wavelength; [8]Price per calibration (10µg) in GBP – 10µg is the minimum required for a calibration using the FPCountR protocol, as dilution of 10µg/1000µl gives 10ng/µl concentration for the highest concentration, which is the recommended minimum for an accurate ECmax quantification.

**Supplementary Table 7. Conversion factors for optical density using microspheres.** Conversion factors (OD/particle) obtained for microsphere calibrations (0.89 µm) in cuvette form and in two plate readers. Conversion factors were converted to particles/OD600 in unist of (particles/ml) per (OD/cm), by normalising for path length of 1 cm, and volume of 1 ml, in order to compare across instruments. Each conversion was obtained from two dilution series of the microspheres, each read in duplicate. Cuvette calibtraions were calculated from two dilution series read once each. The plate reader values are representative of multiple calibration experiments.

**Supplementary Table 8. Relative OD values given by absorbance scans of *E. coli.*** Values from Supplementary Fig. 11A, left panel, showing the OD600-normalised values of relative OD for a range of cell concentrations.

**Supplementary Table 9. Sequences.**

### Supplementary Methods

**Tecan Spark methods.**

**Spark microBCA method**

- Plate
  - Name: COS96ft
  - Plate layout:
  - Plate area: A1-H12
- Protocol
  - Mode: Temperature
  - Temperature control: On
  - Target temperature: 37 [°C]
  - Wait minimum: 36.5 [°C]
  - Wait maximum: 37.5 [°C]
  - .
  - Mode: Wait
  - Wait (Time): 02:00:00 hh:mm:ss
  - Wait position: Incubation
  - .
  - Mode: Temperature
  - Temperature control: On
  - Target temperature: 25 [°C]
  - .
  - Mode: Wait
  - Wait (Time): 00:20:00 hh:mm:ss
  - Wait position: Incubation
  - .
  - Absorbance
  - Name: A562
  - Mode: Absorbance
  - Measurement wavelength: 562 [nm]
  - Number of flashes: 10
  - Settle time: 50 [ms]

**Spark absorbance scan method**

- Plate
  - Name: GRE96ft
  - Plate layout:
  - Plate area: A1-H12
- Protocol
  - Mode: Temperature
  - Temperature control: On
  - Target temperature: 30 [°C]
  - Wait minimum: 29.5 [°C]
  - Wait maximum: 30.5 [°C]
  - .
  - Absorbance Scan
  - Name: Absorbance
  - Wavelength start: 200 [nm]
  - Mode: Absorbance
  - Wavelength end: 1000 [nm]
  - Wavelength step size: 1 [nm]
  - Number of flashes: 1
  - Settle time: 50 [ms]

**Spark fluroescence method**

An example using the red1red1 filter set.

- Plate
  - Name: COS96ft
  - PlateLayout:
  - Plate area: C1-D12;G1-H12
- Protocol
  - Mode: Temperature
  - Temperature control: On
  - Target temperature: 30 [°C]
  - Wait minimum: 29.5 [°C]
  - Wait maximum: 30.5 [°C]

- .
- Fluorescence Intensity
- Name: red1red1040
- Mode: Fluorescence Bottom Reading
- Excitation: Filter
- ExcitationWavelength: 560 [nm]
- ExcitationBandwidth: 20 [nm]
- Emission: Filter
- EmissionWavelength: 620 [nm]
- EmissionBandwidth: 20 [nm]
- Gain: 40 Manual
- Number of flashes: 30
- IntegrationTime: 40 [µs]
- Lag time: 0 [µs]
- SettleTime: 0 [ms]
- Z-Position: 30000 [µm]
- Z-Position mode: Manual
- .
- Fluorescence Intensity
- Name: red1red1050
- Mode: Fluorescence Bottom Reading
- Excitation: Filter
- ExcitationWavelength: 560 [nm]
- ExcitationBandwidth: 20 [nm]
- Emission: Filter
- EmissionWavelength: 620 [nm]
- EmissionBandwidth: 20 [nm]
- Gain: 50 Manual
- Number of flashes: 30
- IntegrationTime: 40 [µs]
- Lag time: 0 [µs]
- SettleTime: 0 [ms]
- Z-Position: 30000 [µm]
- Z-Position mode: Manual
- .
- Fluorescence Intensity
- Name: red1red1060
- Mode: Fluorescence Bottom Reading
- Excitation: Filter
- ExcitationWavelength: 560 [nm]
- ExcitationBandwidth: 20 [nm]
- Emission: Filter
- EmissionWavelength: 620 [nm]
- EmissionBandwidth: 20 [nm]
- Gain: 60 Manual
- Number of flashes: 30
- IntegrationTime: 40 [µs]
- Lag time: 0 [µs]
- SettleTime: 0 [ms]
- Z-Position: 30000 [µm]
- Z-Position mode: Manual
- .
- Fluorescence Intensity
- Name: red1red1070
- Mode: Fluorescence Bottom Reading
- Excitation: Filter
- ExcitationWavelength: 560 [nm]
- ExcitationBandwidth: 20 [nm]
- Emission: Filter
- EmissionWavelength: 620 [nm]
- EmissionBandwidth: 20 [nm]
- Gain: 70 Manual
- Number of flashes: 30
- IntegrationTime: 40 [µs]
- Lag time: 0 [µs]
- SettleTime: 0 [ms]
- Z-Position: 30000 [µm]
- Z-Position mode: Manual
- .
- Fluorescence Intensity
- Name: red1red1080

- Mode: Fluorescence Bottom Reading
- Excitation: Filter
- ExcitationWavelength: 560 [nm]
- ExcitationBandwidth: 20 [nm]
- Emission: Filter
- EmissionWavelength: 620 [nm]
- EmissionBandwidth: 20 [nm]
- Gain: 80 Manual
- Number of flashes: 30
- IntegrationTime: 40 [µs]
- Lag time: 0 [µs]
- SettleTime: 0 [ms]
- Z-Position: 30000 [µm]
- Z-Position mode: Manual
- .
- Fluorescence Intensity
- Name: red1red1090
- Mode: Fluorescence Bottom Reading
- Excitation: Filter
- ExcitationWavelength: 560 [nm]
- ExcitationBandwidth: 20 [nm]
- Emission: Filter
- EmissionWavelength: 620 [nm]
- EmissionBandwidth: 20 [nm]
- Gain: 90 Manual
- Number of flashes: 30
- IntegrationTime: 40 [µs]
- Lag time: 0 [µs]
- SettleTime: 0 [ms]
- Z-Position: 30000 [µm]
- Z-Position mode: Manual
- .
- Fluorescence Intensity
- Name: red1red1100
- Mode: Fluorescence Bottom Reading
- Excitation: Filter
- ExcitationWavelength: 560 [nm]
- ExcitationBandwidth: 20 [nm]
- Emission: Filter
- EmissionWavelength: 620 [nm]
- EmissionBandwidth: 20 [nm]
- Gain: 100 Manual
- Number of flashes: 30
- IntegrationTime: 40 [µs]
- Lag time: 0 [µs]
- SettleTime: 0 [ms]
- Z-Position: 30000 [µm]
- Z-Position mode: Manual
- .
- Fluorescence Intensity
- Name: red1red1110
- Mode: Fluorescence Bottom Reading
- Excitation: Filter
- ExcitationWavelength: 560 [nm]
- ExcitationBandwidth: 20 [nm]
- Emission: Filter
- EmissionWavelength: 620 [nm]
- EmissionBandwidth: 20 [nm]
- Gain: 110 Manual
- Number of flashes: 30
- IntegrationTime: 40 [µs]
- Lag time: 0 [µs]
- SettleTime: 0 [ms]
- Z-Position: 30000 [µm]
- Z-Position mode: Manual
- .
- Fluorescence Intensity
- Name: red1red1120
- Mode: Fluorescence Bottom Reading
- Excitation: Filter
- ExcitationWavelength: 560 [nm]

    - ExcitationBandwidth: 20 [nm]
    - Emission: Filter
    - EmissionWavelength: 620 [nm]
    - EmissionBandwidth: 20 [nm]
    - Gain: 120 Manual
    - Number of flashes: 30
    - IntegrationTime: 40 [µs]
    - Lag time: 0 [µs]
    - SettleTime: 0 [ms]
    - Z-Position: 30000 [µm]
    - Z-Position mode: Manual

**Spark growth curve method**

- Plate
    - Name: COS96ft
    - PlateLayout:
    - Plate area: A1-H12
- Protocol
    - Mode: Temperature
    - Temperature control: On
    - Target temperature: 30 [°C]
    - Wait minimum: 29.5 [°C]
    - Wait maximum: 30.5 [°C]
    - .
    - Kinetic Loop
    - Mode: Kinetic
    - ListOfActionName: Kinetic
    - Kinetic duration: 16:00:00 [hhmmss]
    - Interval time: 00:10:00 [hhmmss]
        - Absorbance
        - Name: OD600
        - Mode: Absorbance
        - Measurement wavelength: 600 [nm]
        - Number of flashes: 10
        - SettleTime: 50 [ms]
        - .
        - Absorbance
        - Name: OD700
        - Mode: Absorbance
        - Measurement wavelength: 700 [nm]
        - Number of flashes: 10
        - SettleTime: 50 [ms]
        - .
        - Fluorescence Intensity
        - Name: green1
        - Mode: Fluorescence Bottom Reading
        - Excitation: Filter
        - ExcitationWavelength: 485 [nm]
        - ExcitationBandwidth: 20 [nm]
        - Emission: Filter
        - EmissionWavelength: 520 [nm]
        - EmissionBandwidth: 10 [nm]
        - Gain: 60 Manual
        - Number of flashes: 30
        - IntegrationTime: 40 [µs]
        - Lag time: 0 [µs]
        - SettleTime: 0 [ms]
        - Z-Position: 30000 [µm]
        - Z-Position mode: Manual
        - .
        - Mode: Shaking
        - Shaking (Double Orbital) Duration: Continuous
        - Shaking (Double Orbital) Position: Incubation
        - Shaking (Double Orbital) Amplitude: 3 [mm]
        - Shaking (Double Orbital) Frequency: 90 [rpm]

**Clariostar methods**

**Excitation spectrum scan – mTagBFP2**

- Basic settings
    - Measurement type: Fluorescence (FI) spectrum

- Microplate name: GREINER 96 F-BOTTOM
- Endpoint settings
  - No. of flashes per well: 20
  - Optic settings
  - Presetname: <user defined settings>
  - No. of wavelength scanpoints: 126
  - Excitation wavelength [nm]: 320.0 -> 445.0; Stepwidth [nm]: 1.0
  - Excitation bandwidth [nm]: 10
  - Emission wavelength [nm]: 475
  - Emission bandwidth [nm]: 16
  - Gain: 1422
  - Gain obtained by: previous gain value (manually entered)
  - Focal height obtained by: focus adjustment performed
  - Focal height [mm]: 5.9
  - Well used for focus adjustment: C1
  - Wavelength used for gain adjust [nm]: 403
- General settings
  - Bottom optic used
  - Spoon type: -
  - Injection needle holder type: -
  - Settling time [s]: 0.5
  - Reading direction: bidirectional, horizontal left to right, top to bottom
  - Target temperature [°C]: 30
  - Target concentration O2 [%]: set off
  - Target concentration CO2 [%]:

**Emission spectrum scan – mTagBFP2**

- Basic settings
  - Measurement type: Fluorescence (FI) spectrum
  - Microplate name: GREINER 96 F-BOTTOM
- Endpoint settings
  - No. of flashes per well: 20
- Optic settings
  - Presetname: <user defined settings>
  - No. of wavelength scanpoints: 136
  - Excitation wavelength [nm]: 390
  - Excitation bandwidth [nm]: 16
  - Emission wavelength [nm]: 420.0 -> 555.0; Stepwidth [nm]: 1.0
  - Emission bandwidth [nm]: 10
  - Gain: 1264
  - Gain obtained by: previous gain value (manually entered)
  - Focal height obtained by: focus adjustment performed
  - Focal height [mm]: 6.1
  - Well used for focus adjustment: C1
  - Wavelength used for gain adjust [nm]: 465
- General settings
  - Bottom optic used
  - Spoon type: -
  - Injection needle holder type: -
  - Settling time [s]: 0.5
  - Reading direction: bidirectional, horizontal left to right, top to bottom
  - Target temperature [°C]: 30
  - Target concentration O2 [%]: set off
  - Target concentration CO2 [%]:

**Excitation spectrum scan – mGFPmut3**

- Basic settings
  - Measurement type: Fluorescence (FI) spectrum
  - Microplate name: GREINER 96 F-BOTTOM
- Endpoint settings
  - No. of flashes per well: 20
- Optic settings
  - Presetname: <user defined settings>
  - No. of wavelength scanpoints: 186
  - Excitation wavelength [nm]: 340.0 -> 525.0; Stepwidth [nm]: 1.0
  - Excitation bandwidth [nm]: 10
  - Emission wavelength [nm]: 555
  - Emission bandwidth [nm]: 16
  - Gain: 1888
  - Gain obtained by: previous gain value (manually entered)
  - Focal height obtained by: previous focal height (focus adjustment performed)

    - Focal height [mm]: 5.4
    - Well used for focus adjustment: E1
    - Wavelength used for gain adjust [nm]: 463
- General settings
    - Bottom optic used
    - Spoon type: -
    - Injection needle holder type: -
    - Settling time [s]: 0.5
    - Reading direction: bidirectional, horizontal left to right, top to bottom
    - Target temperature [°C]: 30
    - Target concentration O2 [%]: set off
    - Target concentration CO2 [%]:

**Emission spectrum scan – mGFPmut3**

- Basic settings
    - Measurement type: Fluorescence (FI) spectrum
    - Microplate name: GREINER 96 F-BOTTOM
- Endpoint settings
    - No. of flashes per well: 20
- Optic settings
    - Presetname: <user defined settings>
    - No. of wavelength scanpoints: 131
    - Excitation wavelength [nm]: 450
    - Excitation bandwidth [nm]: 16
    - Emission wavelength [nm]: 480.0 -> 610.0; Stepwidth [nm]: 1.0
    - Emission bandwidth [nm]: 10
    - Gain: 1456
    - Gain obtained by: previous gain value (manually entered)
    - Focal height obtained by: previous focal height (focus adjustment performed)
    - Focal height [mm]: 5.8
    - Well used for focus adjustment: E1
- General settings
    - Bottom optic used
    - Spoon type: -
    - Injection needle holder type: -
    - Settling time [s]: 0.5
    - Reading direction: bidirectional, horizontal left to right, top to bottom
    - Target temperature [°C]: 30
    - Target concentration O2 [%]: set off
    - Target concentration CO2 [%]:

**Excitation spectrum scan – mCherry**

- Basic settings
    - Measurement type: Fluorescence (FI) spectrum
    - Microplate name: GREINER 96 F-BOTTOM
- Endpoint settings
    - No. of flashes per well: 20
- Optic settings
    - Presetname: <user defined settings>
    - No. of wavelength scanpoints: 301
    - Excitation wavelength [nm]: 320.0 -> 620.0; Stepwidth [nm]: 1.0
    - Excitation bandwidth [nm]: 10
    - Emission wavelength [nm]: 650
    - Emission bandwidth [nm]: 16
    - Gain: 2699
    - Gain obtained by: previous gain value (manually entered)
    - Focal height obtained by: previous focal height (focus adjustment performed)
    - Focal height [mm]: 6.1
    - Well used for focus adjustment: D1
    - Wavelength used for gain adjust [nm]: 520
- General settings
    - Bottom optic used
    - Spoon type: -
    - Injection needle holder type: -
    - Settling time [s]: 0.5
    - Reading direction: bidirectional, horizontal left to right, top to bottom
    - Target temperature [°C]: 30
    - Target concentration O2 [%]: set off
    - Target concentration CO2 [%]:

**Emission spectrum scan – mCherry**

- Basic settings
    - Measurement type: Fluorescence (FI) spectrum
    - Microplate name: GREINER 96 F-BOTTOM
- Endpoint settings
    - No. of flashes per well: 20
- Optic settings
    - Presetname: <user defined settings>
    - No. of wavelength scanpoints: 181
    - Excitation wavelength [nm]: 530
    - Excitation bandwidth [nm]: 16
    - Emission wavelength [nm]: 560.0 -> 740.0; Stepwidth [nm]: 1.0
    - Emission bandwidth [nm]: 10
    - Gain: 1917
    - Gain obtained by: previous gain value (manually entered)
    - Focal height obtained by: previous focal height (focus adjustment performed)
    - Focal height [mm]: 5.8
    - Well used for focus adjustment: C1
    - Wavelength used for gain adjust [nm]: 620
- General settings
    - Bottom optic used
    - Spoon type: -
    - Injection needle holder type: -
    - Settling time [s]: 0.5
    - Reading direction: bidirectional, horizontal left to right, top to bottom
    - Target temperature [°C]: 30
    - Target concentration O2 [%]: set off
    - Target concentration CO2 [%]:

.
